# CHIP protects lysosomes from CLN4 mutant-induced membrane damages

**DOI:** 10.1101/2025.02.18.638932

**Authors:** Juhyung Lee, Jizhong Zou, Wan Nur Atiqah Binti Mazli, Natalie Chin, Michal Jarnik, Layla Saidi, Yue Xu, John Replogle, Michael Ward, Juan Bonifacino, Wei Zheng, Ling Hao, Yihong Ye

## Abstract

Understanding how cells mitigate lysosomal damage is critical for unraveling pathogenic mechanisms of lysosome-related diseases. Here we use organelle-specific proteomics in iPSC-derived neurons (i^3^Neuron) and an *in vitro* lysosome-damaging assay to demonstrate that lysosome damage, caused by the aggregation of Ceroid Lipofuscinosis Neuronal 4 (CLN4)-linked DNAJC5 mutants on lysosomal membranes, serves as a critical pathogenic linchpin in CLN4-associated neurodegeneration. Intriguingly, in non-neuronal cells, a ubiquitin-dependent microautophagy mechanism downregulates CLN4 aggregates to counteract CLN4-associated lysotoxicity. Genome-wide CRISPR screens identify the ubiquitin ligase CHIP as a central microautophagy regulator that confers ubiquitin-dependent lysosome protection. Importantly, CHIP’s lysosome protection function is transferrable, as ectopic CHIP improves lysosomal function in CLN4 i^3^Neurons, and effectively alleviates lipofuscin accumulation and neurodegeneration in a *Drosophila* CLN4 disease model. Our study establishes CHIP-mediated microautophagy as a key organelle damage guardian that preserves lysosome integrity, offering new insights into therapeutic development for CLN4 and other lysosome-related neurodegenerative diseases.

---

Ceroid lipofuscinosis neuronal (CLN) refers to a group of genetically inherited lysosomal storage diseases (LSD) featured with massive neurodegeneration ^1, 2^. These diseases are caused by mutations in a collection of genes (e.g., CLN1, CLN2, CLN3), which lead to lysosomal dysfunction and the buildup of toxic substances in the brain and other tissues ^3, 4^. A common pathologic hallmark of CLN diseases is the accumulation of autofluorescent lipopigments in the form of ceroid or lipofuscin in both neuronal and non-neuronal cells. However, the disease symptoms are mostly manifested in the central nervous system ^5^. CLN, as an uncurable disease, primarily affects children with common progressive symptoms including vision loss, seizures, motor deterioration, cognitive decline, and eventually, profound neurological impairment ^1, 3, 5^. While most CLN cases are autosomal recessive, CLN4 (also known as Kufs disease) is caused by dominant mutations in *DNAJC5* ^6–10^, a gene regulating protein folding and cellular homeostasis.

*DNAJC5*, also named *Cysteine String Protein α* (*CSPα*), encodes a member of the DnaJ/Hsp40 family of molecular chaperones known as DNAJC5. The Hsp40 family members are featured by the presence of a J-domain, which interacts with HSC70 to promote its ATPase cycle ^11, 12^. DNAJC5 also contains a cysteine string domain, which undergoes palmitoylation to facilitate its membrane interactions ^13^. In neurons, DNAJC5 is primarily localized to synaptic vesicles, but it is also found in other membrane compartments such as Golgi-associated vesicles, plasma membrane (PM), and lysosomes ^4^. Owing to its co-chaperone activity ^14^, DNAJC5 plays critical roles in many cellular processes, including calcium regulation ^15^, membrane fusion and exocytosis ^14, 16, 17^, and unconventional secretion of misfolded proteins^18–20^ where it aids in the transport of misfolded cytosolic proteins out of the cell ^21^. DNAJC5 also has a role in microautophagy, a form of autophagy that engulfs smaller portions of the cytoplasm including damaged proteins and organelles by late endosomes/lysosomes ^22^. The diverse roles of DNAJC5 underscore a critical neuroprotective function linked to cellular homeostasis regulation ^23^.

Numerous studies have characterized the biochemical properties of the CLN4-associated DNAJC5 mutants *in vitro* and in cells. It is known that these mutations abolish DNAJC5 palmitoylation ^24^, causing the mutant proteins to be more prone to aggregation ^25–27^. Moreover, these mutations disrupt the association of DNAJC5 with a Golgi-associated compartment ^22^, abolishing its activity in unconventional protein secretion ^20, 22^ while enhancing its association with the lysosomes ^22, 27^. However, it is unclear how these changes disrupt cell homeostasis to cause lipofuscinosis and neuronal cell death.

In this study, we identify lysosomal membrane damage as a key pathogenic mechanism in CLN4 disease. We uncover a ubiquitin- and microautophagy-dependent lysosome protective mechanism that masks CLN4 mutants’ lysotoxicity in non-neuronal cells, which is mediated by the ubiquitin ligase CHIP. Accordingly, CHIP overexpression restores lysosome function in CLN4 mutation-bearing i^3^Neurons and reduces lipofuscin accumulation and neurodegeneration in a *Drosophila* CLN4 disease model. These findings establish CHIP-mediated microautophagy as a promising target of therapeutic intervention in treating lysosome damage-related neurodegenerative diseases.

## Results

### Generation and characterization of human iPSC-derived CLN4 disease models

To model CLN4 disease, we used CRISPR-mediated gene editing to introduce L115R or L116Δ mutation into the endogenous *DNAJC5* locus in a well-characterized iPSC line with verified karyotype and pluripotency (Extended Data Fig. 1a) ^28^. Genomic sequencing confirmed several clones heterozygous (HT) for the L115R or L116Δ allele and a homozygous (HM) L116Δ clone (Extended Data Fig. 1b-d). We selected clones with normal karyotypes and confirmed pluripotency (Extended Data Fig. 1e, f) and transduced them with a lentiviral vector that enabled their differentiation into excitatory neurons (hereafter referred to as i^3^Neurons) via tetracycline-induced expression of neurogenin-2 (NGN2) (Extended Data Fig. 1g) ^29^.

Next, we characterized these cells focusing on phenotypes relevant to the CLN4 disease. CLN4 *DNAJC5* mutants are known to aggregate in an iron-sulfur cluster-, ATP-, and HSC70-dependent manner ^24^. Immunoblotting confirmed that ∼50% of DNAJC5 existed in an SDS-resistant, high molecular weight (HMW) form in cells heterozygous for L115R or L116Δ. In L116Δ HM i^3^Neurons, DNAJC5 was almost entirely present in the HMW form (Fig. 1a, Extended Data Fig. 2a). As expected, substantial amount of the HMW DNAJC5 mutants were present in the NP40-insoluble fractions, consistent with their aggregation properties. While we saw no difference in cell morphology or growth rate between wild-type (WT) and mutant iPSCs, significantly more L115R and L116Δ cells underwent apoptosis compared to WT cells after 16 days (d16) in differentiation, and L116Δ HM cells exhibited the most severe phenotype (Fig. 1b, c).

**Fig. 1:**
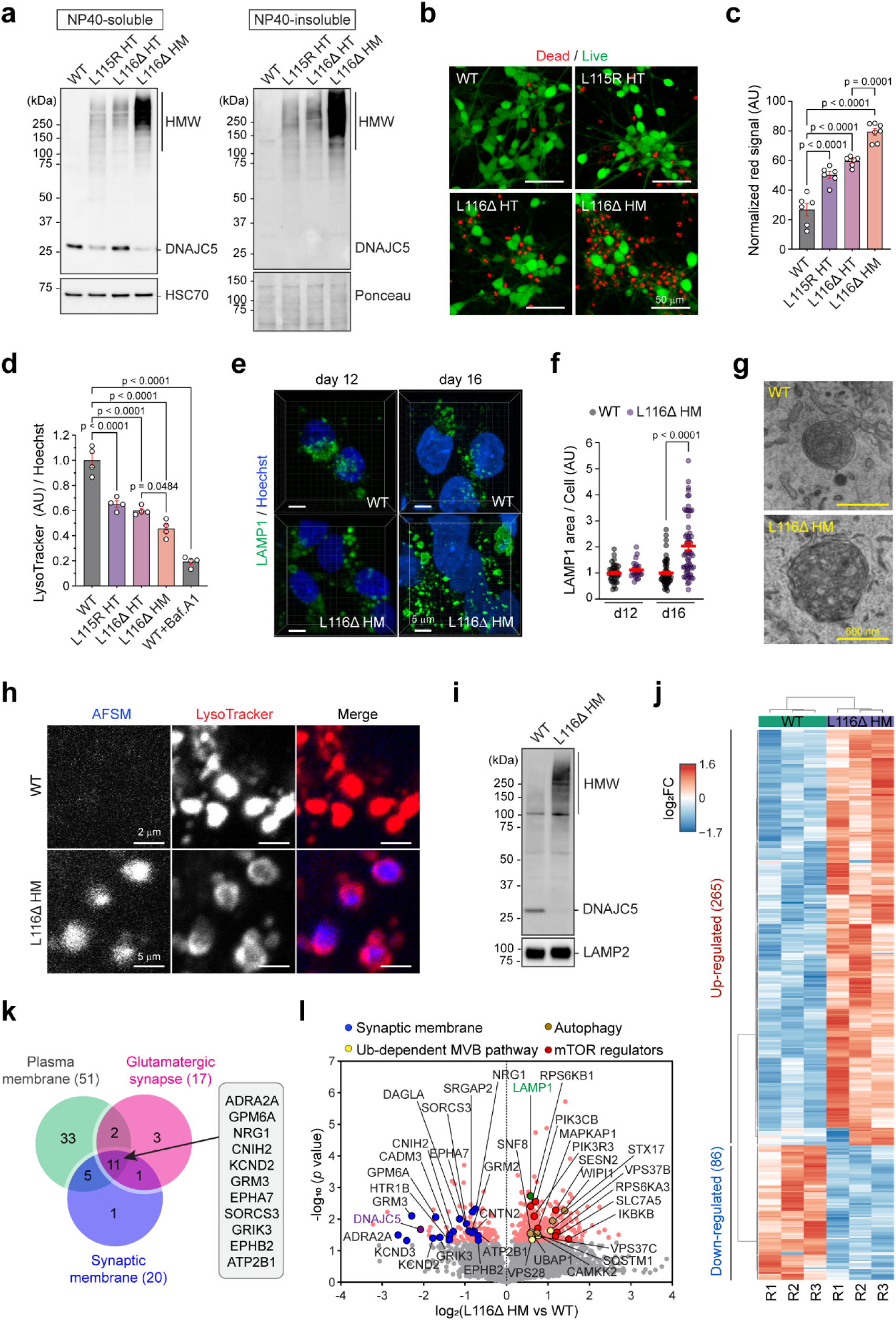
Disrupted lysosome homeostasis in iPSC-derived i^3^Neuron bearing CLN4 mutations. **a**, Representative immunoblots of DNAJC5 in i^3^Neurons of the indicated genotypes at day 16 in differentiation (d16). The blots represent 4 independent experiments. HMW, high molecular weight species, WT, wild-type; HT, heterozygous; HM, homozygous. **b,** i^3^Neurons of the indicated genotypes at d16 were stained by green-fluorescent calcein-AM to indicate intracellular esterase activity (live cells) and red-fluorescent ethidium homodimer-1 to indicate loss of plasma membrane integrity (dead cells). Scale bars, 50 µm. **c,** Quantification of the experiments in b. Each dot represents a randomly selected field. P-values were determined by one-way ANOVA. n=2 biological repeats. **d,** i^3^Neurons of the indicated genotypes at d16 were stained with LysoTracker Red (0.2 µM) and Hoechst33442 (1 µg/mL). The fluorescence intensities were measured by a plate reader. Cells treated with Bafilomycin A1 (Baf.A1) serve as a positive control. AU, arbitrary unit. P-values were determined by one-way ANOVA; n=4 biological repeats. **e,** WT and L116Δ homozygous (HM) i^3^Neurons at the indicated differentiation stage were stained by anti-LAMP1 antibodies (green) and Hoechst (blue). Scale bars, 5 µm. **f,** Quantification of LAMP1-positive area in randomly selected cells (n>20) in three independent experiments represented by e. P-value by unpaired Student’s t-test. **g,** Representative EM images of lysosomes in WT and L116Δ HM i^3^Neurons at d16. **h,** WT and L116Δ HM i^3^Neurons at d16 were stained by LysoTracker Red and imaged. **i,** A fraction of DNAJC5 positive membrane fractions from the indicated i^3^Neurons were analyzed by immunoblotting. **j,** A heat map shows the proteins up- or down-regulated on the DNAJC5-positive organelles in L116Δ HM i^3^Neurons, as determined by three mass spectrometry analyses. **k,** A Venn diagram shows the three major categories of the proteins down-regulated on the DNAJC5-positive organelles by L116Δ HM mutation. **l,** A volcano plot highlighting the proteins up- and down-regulated on the DNAJC5-positive organelles by L116Δ HM mutation. Error bars in c, d, f represent mean ± standard error of the mean (s.e.m.).

Since CLN4 is an LSD, we characterized the lysosomes by LysoTracker staining, which preferentially labels acidic lysosomes. Fluorescence intensity measurement indicated a significant reduction of LysoTracker signal in mutant cells compared to the WT control, with L116Δ HM i^3^Neurons showing the most pronounced decrease (Fig. 1d). Interestingly, while imaging live cells stained with LysoTracker confirmed the pH increase phenotype in d16 L116Δ HM i^3^Neurons, no significant change in lysosomal pH was observed in d12 L116Δ HM i^3^Neurons (Extended Data Fig. 2b). Staining with Magic Red, a fluorogenic dye indicative of the lysosomal protease Cathepsin B (CTSB) activity, showed reduced CTSB activity in mutant cells, mirroring the LysoTracker result (Extended Data Fig. 2c). These findings highlight lysosomal dysfunction as a primary defect dose-dependently induced by CLN4 mutant alleles, like via a yet-to-be defined gain-of-toxic activity suggested previously ^22, 27, 30^.

To further investigate the molecular basis of CLN4 mutant-induced neurotoxicity, we focused on L116Δ HM i^3^Neurons due to their severe phenotype. Immunostaining for LAMP1, a lysosomal membrane protein, in d16 i^3^Neurons showed increased LAMP1 signal in L116Δ HM i^3^Neurons compared to WT control (Fig. 1e, right panels); L116Δ HM i^3^Neurons also contained many enlarged, irregularly shaped lysosomes. Interestingly, these abnormalities were not obvious at d12 (Fig. 1e, f). Transmission electron microscopy (TEM) revealed spherical lysosomes with dense core of 0.1–1 µm in diameter in WT i^3^Neurons, while in L116Δ HM cells, we frequently detected giant lysosomes (2–3 µm in diameter) containing undigested materials. These enlarged lysosomes often had rough or ruptured membranes (Fig. 1g, Extended Data Fig. 2d). These observations, together with the reduced LysoTracker staining and CTSB activity, suggest that lysosomes might be damaged in CLN4 mutant i^3^Neurons.

A hallmark of CLN diseases is the accumulation of autofluorescent storage material (AFSM). Indeed, confocal microscopy revealed many autofluorescent puncta in L116Δ HM i^3^Neurons but not WT cells (Fig. 1h). Most AFSM puncta were colocalized with LysoTracker signals, indicating lysosome as their source of origin.

To further elucidate the organelle homeostasis defects in L116Δ i^3^Neurons, we performed an organelle-based proteomics study. We isolated microsomes from WT and L116Δ HM i^3^Neurons using gradient centrifugation and analyzed DNAJC5-enriched membrane fractions by mass spectrometry (Fig. 1i, Extended Data Fig. 2e, f). Based on the known subcellular localizations of DNAJC5 ^4^, these DNAJC5-positive membranes should include lysosomes, synaptic vesicles, Golgi-associated vesicles, and the PM. Mass spectrometry analysis identified 265 proteins up-regulated by at least 1.5-fold and 86 proteins down-regulated similarly on membranes of L116Δ HM i^3^Neurons (Fig. 1j, Supplementary Table 1). Gene Ontology (GO) pathway analysis showed that downregulated proteins were predominantly synaptic and PM proteins linked to synaptic transmission (Fig. 1k). By contrast, up-regulated proteins were mostly involved in pathways associated with lysosomal homeostasis including ubiquitin-dependent microautophagy (also named multivesicular body or MVB) and mTOR signaling (Fig. 1l, Extended Data Fig. 2g). These results suggest that CLN4 mutants disrupt synaptic membrane proteome and lysosome homeostasis.

### CLN4-associated DNAJC5 aggregates damage lysosomal membranes

Several lines of evidence suggested that the observed lysosomal defects in CLN4 i^3^Neurons likely resulted from membrane destabilization rather than the inhibition of the vacuolar ATPase (v-ATPase) complex. First, while co-immunoprecipitation readily detected an interaction between WT DNAJC5 and endogenous ATP6V1G2, a cytoplasmic subunit of the v-ATPase complex, as shown previously ^31^, the interaction of ATP6V1G2 with CLN4 mutants was barely detectable (Extended data Fig. 3a), even though the CLN4 mutants are known to associate with the lysosomes better than WT DNAJC5 ^22^. Thus, it seemed unlikely that CLN4 mutants could co-aggregate with the v-ATPase complex. Importantly, electron microscopy revealed many abnormal lysosomes with disrupted membranes in L116Δ HM i^3^Neurons (Extended data Fig. 2d), consistent with the lysosomal recruitment of ESCRT components, a phenotype known as a cellular stress response to lysosome damages ^32–36^.

**Fig. 2:**
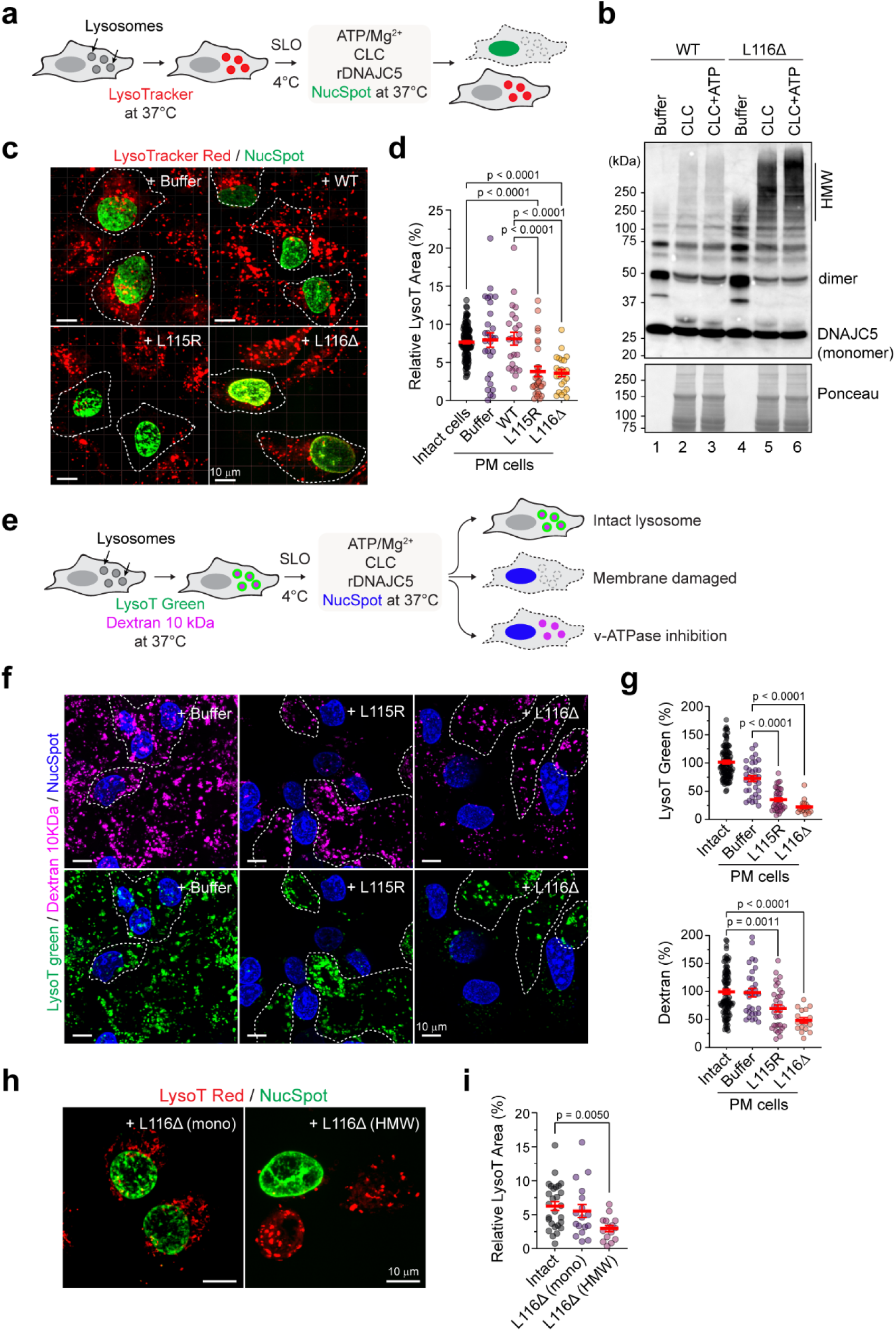
CLN4 mutant aggregates can damage lysosomal membranes. **a,** A schematic diagram showing the *in vitro* lysosome-damaging assay. SLO, Streptolysin O, CLC; calf liver cytosol. **b,** Purified WT DNAJC5 or L116Δ mutant incubated with buffer or CLC with or without ATP at 37 °C for 30 min were analyzed by immunoblotting. HMW, high molecular weight species. **c,** U2OS cells stained with LysoTracker Red (LysoT Red) were treated with SLO and then incubated with either buffer or the indicated DNAJC5 proteins with CLC, ATP and a cell impermeable dye NucSpot green at 37 °C for 30 min. Cells were imaged immediately. Scale bars, 10 µm. **d,** Quantification of the relative LysoT-positive areas in individual permeabilized (PM) cells treated with buffer or the indicated DNAJC5 proteins. Intact cells as indicated by the lack of NucSpot staining serves as a reference. P-values were determined by one-way ANOVA from three biological repeats. **e,** The modified lysosome damaging assay using Dextran (10 kDa)-loaded cells stained with LysoTracker green. **f,** U2OS cells loaded with Dextran (10 kDa, magenta) were stained with LysoTracker green (green) and then treated with SLO. Permeabilized cells were then treated CLC with ATP and the indicated CLN4 mutant proteins or buffer as a negative control. Dashed lines indicate intact cells. Scale bars, 10 µm. **g,** Quantification of the relative LysoT positive areas (top panel) or Dextran-positive areas (bottom panel) in individual permeabilized (PM) cells treated with buffer or the indicated DNAJC5 proteins plus ATP and CLC. P-values were determined by one-way ANOVA from two independent experiments. **h,** As in c, excepted that permeabilized cells were treated with monomeric L116Δ (left) or purified L116Δ HMW species (right) plus ATP but in the absence of cytosol. Scale bars, 10 µm. **i,** Quantification of the relative LysoT-positive areas in experiment h. P-values were determined by unpaired student’s t-test from two biological repeats. Error bars in d and g represent mean ± standard error of the mean (s.e.m.). Confocal images in c, f, and h were processed using maximum intensity projection to represent the fluorescence signal across the z-stacks.

**Fig. 3:**
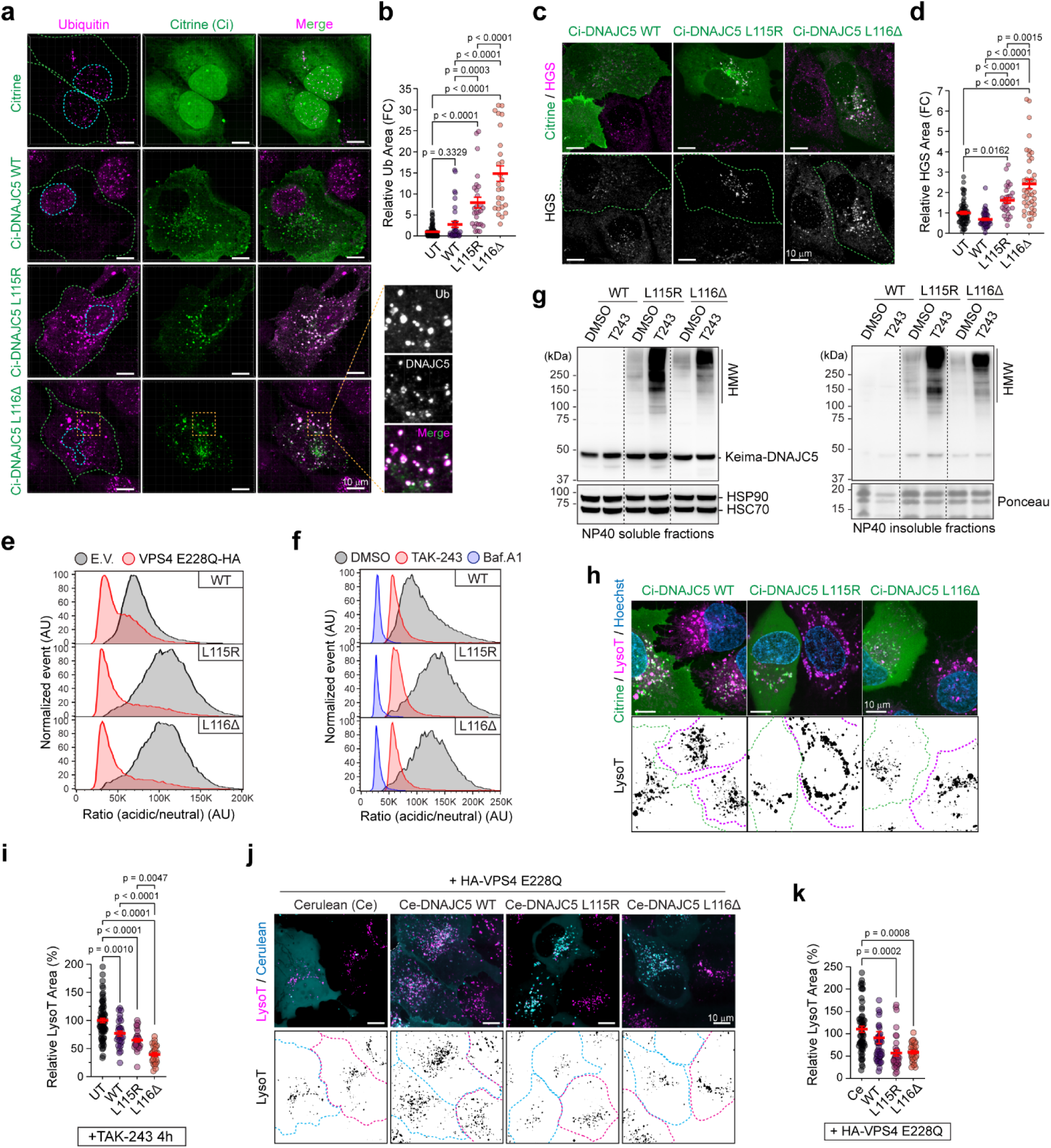
A ubiquitin- and ESCRT-dependent mechanism counteracts CLN4-induced lysotoxicity in non-neuronal cells. **a,** U2OS cells transfected mCitrine (Ci) or Ci-tagged DNAJC5 variants were methanol-fixed and stained with ubiquitin antibody (FK2) (magenta). The box-indicated area in the bottom panels is enlarged to show the partial co-localization of DNAJC5 with ubiquitin (Ub) (right panels). Green and cyan-dotted lines highlight cell boundaries and nuclei of transfected cells, respectively. Scale bars, 10 µm. **b,** Quantification of the relative cytoplasmic Ub puncta areas in cells transfected with the indicated DNAJC5 variants. Untransfected (UT) cells serve as a reference. P-values were determined by one-way ANOVA from two biological repeats. **c,** As in a, except that cells transfected with the indicated DNAJC5 variants were stained with antibodies against HGS. Scale bars, 10 µm. **d,** Quantification of the relative cytoplasmic HGS puncta areas as shown in c. P-values were determined by one-way ANOVA from two biological repeats. **e,** HEK293T cells stably expressing Keima-tagged WT DNAJC5 or the indicated CLN4 mutants were transfected with a dominant negative form of VPS4 (E228Q) and analyzed by flow cytometry. **f,** HEK293T cells expressing Keima-tagged DNAJC5 variants were treated with Baf.A1 (100 nM) or TAK-243 (1 µM) for 4 h and then analyzed by flow cytometry. **g,** HEK293T cells expressing Keima-tagged DNAJC5 variants were treated with DMSO as a control or TAK-243 (T243, 1 µM) for 16 h. NP40-soluble and -insoluble fractions were analyzed by immunoblotting. **h,** U2OS cells transfected with Ci-tagged DNAJC5 variants were treated with TAK-243 (1 µM, 4 h) and stained with LysoTrack Red (magenta) and Hoechst. Scale bars, 10 µm. **i,** Quantification of the relative LysoT-positive areas in transfected cells after 4 h treatment with TAK-243 (1 µM). Untransfected cells serve as internal negative controls. P-values from unpaired student’s t-test (left) or one-way ANOVA (right) from two biological repeats. **j,** As in h, except that cells transfected with Celurean (Ce) or Ce-tagged DNAJC5 variants together with VPS4 E228Q-HA were stained with LysoTracker Red (magenta). Scale bars, 10 µm. **k,** Quantification of the relative LysoT positive areas in cells expressing the indicated proteins. P-values are determined by one-way ANOVA from two biological repeats. Error bars in b, d, i and k represent mean ± standard error of the mean (s.e.m.). In a, c, h, and j, z-stack images were processed using maximum intensity projection.

We postulated that lysosome-associated CLN4 aggregates might have a lysosome damaging activity. To test this idea, we developed an *in vitro* assay using semi-permeabilized U2OS cells (Fig. 2a). We labeled functional lysosomes with a LysoTracker dye and then permeabilized the PM in a fraction of the cells with the pore forming toxin streptolysin O (SLO). After PM permeabilization and removal of the cytosol, cells were incubated with recombinant DNAJC5 together with a membrane impermeable dye (to label permeabilized cells), ATP, and concentrated cow live cytosol. We included cytosol in this experiment because immunoblotting demonstrated that incubation with ATP and cytosol caused CLN4 mutants to form HMW aggregates (Fig. 2b), analogous to those observed in cells. Confocal microscopy showed that addition of L115R or L116Δ to SLO-treated cells reduced LysoTracker signal specifically in permeabilized cells, whereas in buffer or WT DNAJC5-treated cells, permeabilized and unpermeabilized cells had similar LysoTracker signal (Fig 2c, d). These results further suggested that CLN4 DNAJC5 mutants can directly impair lysosomes.

To conclusively demonstrate the membrane destabilizing activity of the CLN4 mutants, we repeated the lysosome-damaging experiment after loading LysoTracker green-stained cells with Alexa^568^-labeled low molecular weight Dextran (Fig. 2e). When these cells were treated with the lysosome-damaging compound LLOMe, both LysoTracker and Dextran signals were lost due to membrane damage (Extended Data Fig. 3b). On the other hand, when SLO-permeabilized cells were incubated in a buffer without ATP, v-ATPase was immediately inhibited but lysosomes remained intact. Consequently, we observed a rapid reduction of the LysoTracker signal, but the Dextran signal remained unaffected (Extended Data Fig. 3c, middle panels). As expected, the addition of ATP maintained the v-ATPase activity, allowing cells to retain both LysoTracker and Dextran signals (bottom panels). Thus, this sensitive assay is capable of distinguishing membrane damage from v-ATPase inhibition. We then incubated SLO-treated cells preloaded with Dextran and LysoTracker with cytosol, ATP, and CLN4 mutants, and observed reduced LysoTracker and Dextran signals in CLN4 mutant-treated cells compared to mock-treated cells (buffer) (Fig. 2f, g). These results confirmed that both L115R and L116Δ mutants can destabilize lysosomal membrane to cause membrane leakage.

To see whether it is the aggregated CLN4 species that induces lysotoxicity, we first incubated purified L116Δ with cow liver cytosol and ATP, and then isolated the HMW species by centrifugation through a sucrose cushion. We incubated SLO-permeabilized cells with either untreated L116Δ (mostly monomer and dimer) or purified L116Δ HMW species in the absence of cytosol. Under this condition, only HMW L116Δ could reduce LysoTracker signal (Fig. 2h, i), suggesting that L116Δ aggregates are responsible for the lysotoxic activity, and the role of the cytosol is to promote CLN4 aggregation. Since lysosomal defects were only seen in mature CLN4 neurons but not in immature neurons (Fig. 1e, f) or U2OS cells overexpressing L116Δ (Extended Data Fig. 4a), it seems that non-neuronal cells have evolved a mechanism to counteract CLN4 mutants’ lysotoxicity.

### Ubiquitin-dependent microautophagy safeguards lysosomes from CLN4-induced membrane damages in non-neuronal cells

The recruitment of the ESCRT components to lysosomes in L116Δ HM i^3^Neurons prompted us to investigate whether CLN4 mutants could activate ubiquitin-dependent microautophagy in non-neuronal cells. Indeed, immunostaining detected many ubiquitin-positive cytoplasmic puncta in U2OS cells expressing mCitrine (Ci)-tagged L115R or L116Δ; most puncta also contained DNAJC5, revealing their identity as lysosomes (Fig. 3a, b). By contrast, few ubiquitin puncta were observed in cells expressing WT Ci-DNAJC5 or mCitrine. Moreover, the ubiquitin-binding component of the ESCRT0 complex hepatocyte growth factor-regulated tyrosine kinase substrate (HGS) was also recruited to CLN4-positive puncta (Fig. 3c, d). Denatured immunoprecipitation demonstrated that membrane-associated CLN4 mutants were more ubiquitinated than WT DNAJC5 in non-neuronal cells overexpressing these proteins (Extended Data Fig. 4b). Together, these results suggest that CLN4 mutants induce ubiquitin build-up on lysosomes with some conjugates attached to the mutant proteins. Lysosome-associated ubiquitination then recruits HGS and activates microautophagy.

To see whether CLN4 mutants are microautophagy substrates, we expressed Keima-tagged DNAJC5 variants in HEK293T cells. Keima is a pH sensitive green fluorescence protein that displays distinct fluorescent spectrum in different pH environments ^37^. Flow cytometry showed that CLN4 mutants had increased lysosomal translocation activity compared to WT DNAJC5, but like WT DNAJC5, their lysosomal translocation was inhibited by an ATPase inactive VPS4 mutant (VPS4 E228Q) that blocks ESCRT-dependent microautophagy, and by the ubiquitin E1 inhibitor TAK-243 (Fig. 3e, f). Immunoblotting confirmed that in cells treated with TAK-243, CLN4 mutants were accumulated in HMW form (Fig. 3g). Together, these results suggest that CLN4 mutants are targeted for degradation by ubiquitin- and ESCRT-dependent microautophagy in non-neuronal cells.

In addition to ESCRT-dependent microautophagy, mammalian cells also use ESCRT-independent microautophagy to degrade abnormal cytosolic proteins ^38^. To test whether ESCRT-independent microautophagy also targets CLN4 mutants, we treated cells expressing CLN4 mutants with GW4869. GW4869 is a neutral sphingomyelinase inhibitor that blocks ceramide synthesis, which is required for ESCRT-independent microautophagy ^38^. Like TAK-243, GW4869 treatment also reduced the lysosomal translocation of CLN4 mutants, causing these proteins to accumulate in cytoplasmic aggregates (Extended Data Fig. 4c, d).

To test whether microautophagy counteracts CLN4 mutants’ lysotoxicity in non-neuronal cells, we treated U2OS cells transfected with the CLN4 mutants with TAK-243 to inhibit microautophagy. Unlike in untreated cells, we now detected a reduction in LysoTracker signal in L115R- or L116Δ-expressing cells, and to a lesser extent, also in WT DNAJC5-expressing cells compared to untransfected cells (Fig. 3h, i). As expected, cells co-expressing VPS4 E228Q with CLN4 mutants also had reduced LysoTracker signal (Fig. 3j, k), so were the DNAJC5 L116Δ-expressing cells exposed to GW4869 treatment (Extended Data Fig. 4e, f). Thus, both ESCRT-dependent and ESCRT-independent microautophagy can target CLN4 mutants, ameliorating lysosome dysfunction in non-neuronal cells expressing these mutants.

### CRISPR screens identify CHIP-mediated microautophagy as a lysosome guardian in non-neuronal cells

To identify the factors conferring ubiquitin-dependent lysosome protection, we conducted two CRISPR screens using HEK293T cells stably expressing Keima-WT DNAJC5 and a Keima-tagged DNAJC5 mutant lacking the HSC70 binding J domain (ΔJ). These proteins undergo microautophagy similarly as CLN4 mutants ^22^, but are less toxic, and therefore, more suitable for long-term overexpression. We infected Keima-DNAJC5 and Keima-DNAJC5 ΔJ mutant cells with a lentiviral library targeting the ∼20,000 human genes each with six sgRNAs (Extended Data, Fig. 5a) ^39^. We then used FACS to isolate cells with increased neutral-to-acidic Keima ratio, indicative of impaired microautophagy. High-throughput sequencing in two biological repeats for each screen identified sgRNAs and the corresponding target genes enriched in cells with defective microautophagy (Fig. 4a). Among them, 85 were deemed as high-confident hits because they were statistically significant in both WT and ΔJ screens (Fig. 4b). Moreover, many hits in this list are known microautophagy regulators such as components of the ESCRT complexes (Fig. 4c). We also identified ATP6V0C, an integral membrane component of the v-ATPase complex, whose inactivation is expected to deacidify the lysosomes. Intriguingly, a ubiquitin ligase named CHIP/STUB1 and its cognate conjugating enzyme UBE2N were also among the high confident list, and small hairpin RNA (shRNA)-mediated knockdown validated them as positive regulators of DNAJC5 microautophagy (Fig. 4d). We decided to focus our studies on CHIP because it was annotated by BioGRID as a component of the DNAJC5 interactome (Fig. 4b) ^31^. Indeed, co-immunoprecipitation confirmed that CHIP could bind both WT DNAJC5 and the CLN4 mutants independent of the J domain (Fig. 4e).

**Fig. 4:**
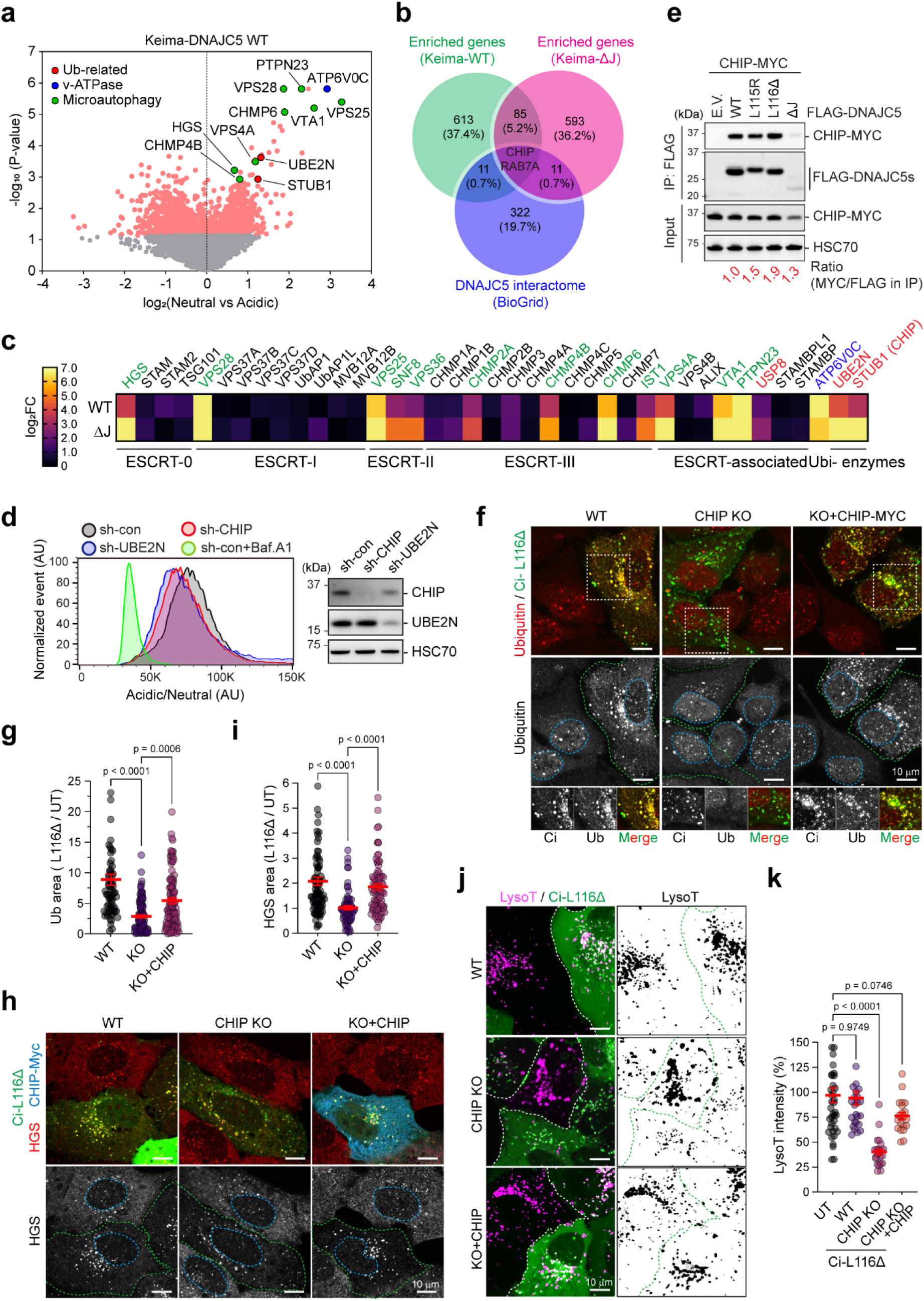
CHIP-mediated ubiquitination protects lysosomes from DNAJC5 L116Δ-induced lysosome damage. **a,** A volcano plot shows the selected sgRNAs enriched in the neutral population from the Keima-DNAJC5 WT-based CRISPR screen. **b,** A Venn diagram shows CHIP and RAB7A as the only overlapped genes from the two CRISPR screens, which were also identified in the DNAJC5-interactome. **c,** A heat map shows the relative significance of the positive hits from the CRISPR screens. **d,** HEK293T cells expressing Keima-DNAKC5 WT were transfected with shRNA-expressing constructs targeting the indicated genes and analyzed by flow cytometry. Con, Control. **e,** HEK293T cells were transfected with FLAG-tagged DNAJC5 variants as indicated or an empty vector (EV) together with MYC-tagged CHIP. FLAG beads were used to pull down DNAJC5. Bound proteins were analyzed by immunoblotting together with a fraction of the input samples. **f,** WT or CHIP knockout (KO) cells were transfected with Ci-tagged L116Δ, methanol-fixed, and stained with ubiquitin antibodies. In the right panels, CHIP KO cells were co-transfected with Ci-L116Δ and CHIP-MYC. Green- and blue-dotted lines indicate transfected cells and nuclei respectively. Note that CHIP expression rescues the defect in lysosome-associated ubiquitination in CHIP KO cells. Scale bars, 10 µm. **g,** Quantification of the positive cytoplasmic ubiquitin area normalized by that in untransfected (UT) cells in f. n= 2 biological repeats. **h,** As in f, excepted that cells were stained with HGS and MYC antibodies. Scale bars, 10 µm. **i,** Quantification of h as in g. P-values in g and I are determined by one-way ANOVA from two biological repeats. **j,** WT and CHIP KO U2OS cells were transfected with stained with Ci-L116Δ, stained with LysoTracker Red (magenta) and imaged. Where indicated, CHIP KO cells were co-transfected with Ci-L116Δ and CHIP-MYC. Note that only in CHIP KO background, L116Δ expression reduced LysoTracker signal. Scale bars, 10 µm. **k,** Quantification of the relative LysoTracker signal in j. P-values are determined by one-way ANOVA from two biological repeats. In f, h, and j, z-stack images were processed using maximum intensity projection.

**Fig. 5:**
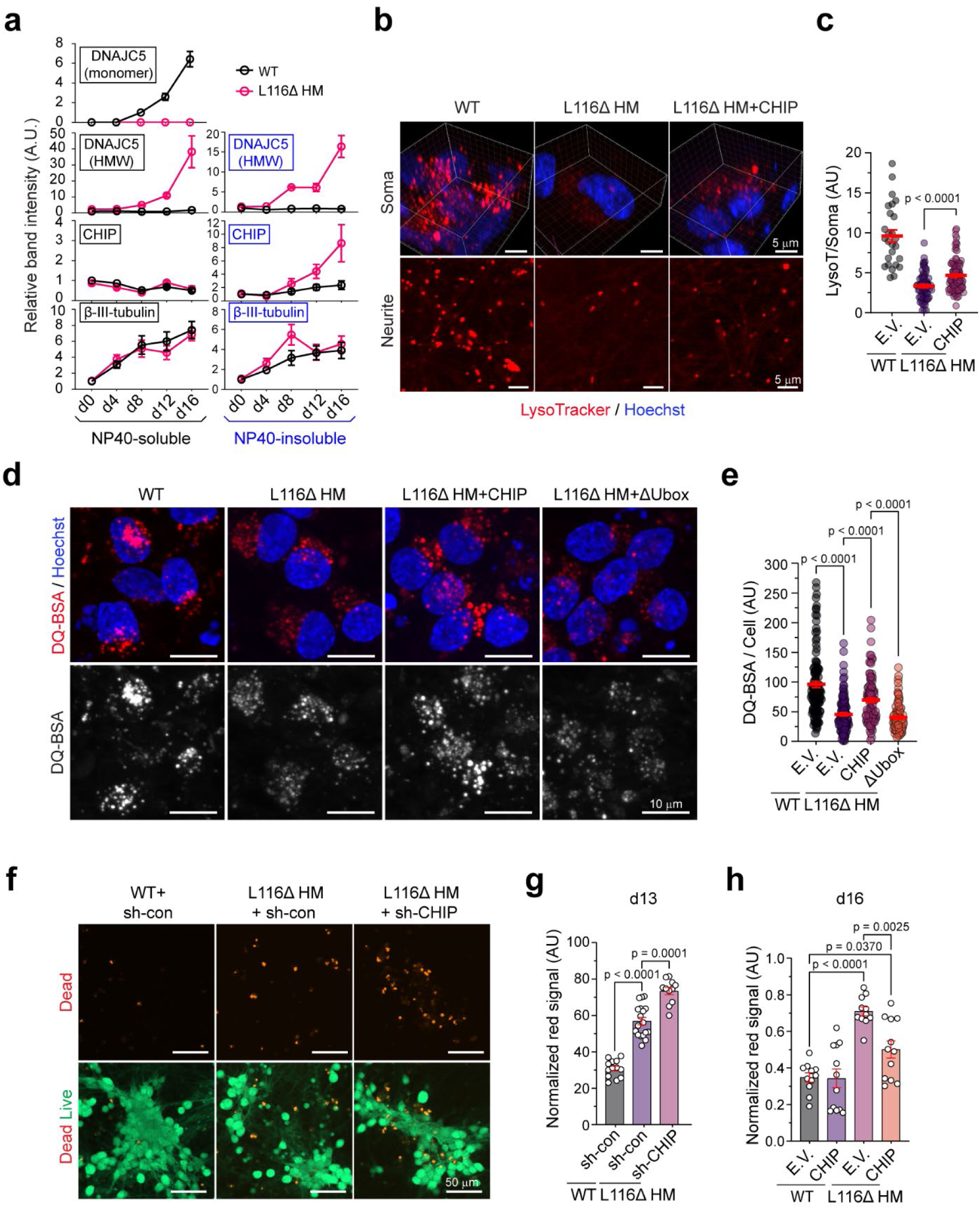
CHIP expression partially rescues the lysosomal defect and cell death phenotypes in L116Δ i^3^Neurons. **a,** Quantification of the levels of the indicated proteins in NP40-soluble or insoluble fractions during i^3^Neuron differentiation. Representative blots are shown in Extended data Fig. 6a. Error bars represent s.e.m. of three biological repeats. **b,** WT and L116Δ HM i^3^Neurons were mock infected (left and middle panels) or infected with CHIP-expressing lentiviruses at d6. Cells were stained with LysoTracker Red and Hoechst at d16 and imaged. Shown are 3D reconstructed view (top panels) of soma or a z section of a neurite-enriched region (bottom panels). Scale bars, 5 µm. **c,** Quantification of LysoT signal in soma as shown in b. P-value is by unpaired student’s t-test. Error bars represent s.e.m. of individual cells (n>20) from two biological repeats. **d**, As in b except that L116Δ HM i^3^Neurons were also infect with a lentivirus expressing CHIP ΔU-box for comparison and that cells at d16 were treated with DQ-BSA (40 µg/ml) for 6 h and then stained with Hoechst. Z-stack images were processed using maximum intensity projection. Scale bars, 10 µm. **e,** Quantification of the relative DQ-BSA fluorescence intensity as shown in d. Error bars represent s.e.m of individual cells from two biological repeats. P-values are determined by one-way ANOVA. **f,** WT or L116Δ HM i^3^Neurons were infected with lentiviruses expressing either control (sh-con) or CHIP specific shRNA. Cells at d13 were stained by green calcein-AM and red ethidium homodimer-1. Scale bars, 50 µm. **g,** Quantification of cell death as shown in f from two biological repeats. Each dot represents a randomly selected field. Error bars represent means ± s.e.m. P-value are determined by one-way ANOVA. **h,** WT or L116Δ HM i^3^Neurons were infected with lentiviruses carrying either an empty vector or a CHIP-expressing cassette. Cell death at d16 were measured as in f. Shown is the quantification results as in g. Error bars represent means ± s.e.m. of two biological repeats. Each data point represents a randomly selected field. P-value are determined by one-way ANOVA.

CHIP (C-terminus of HSC70-Interacting Protein) is a U-box containing ubiquitin ligase that interacts with HSP90 and HSC70 via several TRP motifs. Mutations in CHIP have been widely linked to neurodegenerative diseases, but the underlying mechanisms are unknown ^40, 41^. To better characterize the role of CHIP in microautophagy, we used CRISPR to generate CHIP knockout (KO) cells. As a positive control, we also generated CHMP6 (a component of ESCRT-III) KO cells. Depletion of either CHMP6 or CHIP significantly reduced lysosomal translocation of Keima-DNAJC5 (Extended Data Fig. 5b). Noticeably, the lysosomal translocation of Keima-tagged CLN4 mutants was also diminished in CHIP KO cells (Extended Data Fig. 5c), further confirming its role in microautophagy.

To further characterize the role of CHIP in microautophagy, we used confocal microscopy to test whether CHIP is required for lysosomal accumulation of ubiquitin and HGS in U2OS cells expressing CLN4 mutants using L116Δ as a representative. Immunostaining showed that unlike WT cells, the lysosomal accumulation of ubiquitin and HGS in L116Δ-expressing CHIP KO cells were not obvious, but re-expressing CHIP rescued this phenotype (Fig. 4f-i). Furthermore, denatured immunoprecipitations showed that CHIP depletion reduced the ubiquitination of both WT and the L116Δ DNAJC5 mutant (Extended data Figure 5d, e). Thus, CHIP promotes DNAJC5 ubiquitination, particularly those associated with CLN4 mutants, resulting in HGS recruitment to lysosomes.

To see whether CHIP-mediated ubiquitination counteracts CLN4-mutants’ lysotoxicity, we transfected WT or CHIP KO U2OS cells with Ci-L116Δ and then stained these cells with LysoTracker. Like in E1 inhibitor-treated cells, Ci-L116Δ expression in CHIP KO cells also destabilized lysosomes to reduce LysoTracker staining. This phenotype was rescued when WT CHIP was co-transfected (Fig. 4j, k). Together, these results suggest that in non-neuronal cells, CHIP stimulates ubiquitination on CLN4-containing lysosomes, activating microautophagy to counteract the membrane damaging activity of the CLN4 mutants.

### CHIP rescues lysosomal defects and cell death in L116Δ i^3^Neurons

What accounts for the differential sensitivity to CLN4-mediated lysotoxicity between immature and mature i^3^Neurons? To address this question, we used immunoblotting to compare the expression of CHIP and DNAJC5 during the differentiation of WT and L116Δ HM i^3^Neurons. Our data suggested that DNAJC5 expression was increased after d8 in both WT and L116Δ HM cells, and as expected, L116Δ was mostly in the HMW form (Fig. 5a, Extended Data Fig. 6a). By contrast, CHIP protein solubilized by the detergent NP40 was largely unchanged throughout the differentiation and between WT and L116Δ cells. However, we observed an increase of CHIP in NP40-insoluble fractions, starting at d8 and more pronounced in L116Δ cells. The accumulation of CHIP in NP40-insoluble fractions might reflect a change in its activity because this phenotype correlated with the accumulation of HMW L116Δ aggregates (Fig. 5a, Extended Data Fig. 6a). Moreover, transducing i^3^Neurons with Keima-DNAJC5-expressing lentiviruses revealed a modest reduction of DNAJC5 lysosomal translocation after d12, suggesting reduced microautophagy (Extended Data Fig. 6b, c). Collectively, these results suggested a reduction in CHIP activity during late stage of L116Δ i^3^Neuron differentiation, which diminishes microautophagy and causes L116Δ aggregate to accumulate. These changes explain why significant lysosome defects and cell death were only noticed in L116Δ HM cells after d12 in differentiation.

To further test the role of microautophagy in neuroprotection, we asked whether ectopically expressing CHIP in i^3^Neuron could alleviate lysosomal damage caused by endogenous L116Δ. To avoid overexpression artifact, we used a neuron-specific synapsin promoter activated after d8 in differentiation to drive CHIP expression in i^3^Neurons. Immunoblotting revealed a modest, but reproducible reduction of total L116Δ aggregate in L116Δ HM cells expressing CHIP compared to cells with no ectopic CHIP (Extended Data Fig. 6d, e). This phenotype was more pronounced when affinity-purified lysosomes from control and CHIP-expressing L116Δ HM i^3^Neurons were analyzed (Extended Data Fig. 6f).

As anticipated, CHIP-expressing L116Δ i^3^Neurons had increased LysoTracker signal in both soma and neurites (Fig. 5b, c). Consistent with improved acidity, DQ-BSA-based lysosomal activity assay showed partially restored hydrolase activity in L116Δ HM i^3^Neurons by WT but not the ΔUbox CHIP mutant lacking the U-box (Fig. 5d, e). Importantly, CHIP knockdown in L116Δ-overexpressing i^3^Neurons enhanced L116Δ-induced cell death (Fig. 5f, g), whereas ectopic expression of CHIP in L116Δ HM i^3^Neurons reduced differentiation-associated cell death (Fig. 5h). Collectively, these results demonstrated that ectopic CHIP improves lysosome homeostasis and mitigates cell death in L116Δ HM i^3^Neurons.

### CHIP rescues lipofuscin accumulation and neurodegeneration in a *Drosophila* CLN4 model

To see whether CHIP could rescue CLN4 mutant-associated disease phenotypes *in vivo*, we tested its function in a recently established *Drosophila* CLN4 disease model ^27^. We first generated a transgenic line carrying Keima-*Csp1*, the *Drosophila* homolog of DNAJC5 (dDNAJC5). We also generated transgenic flies bearing *Drosophila* WT CHIP (dCHIP) or dCHIP lacking the TRP or U-box-coding sequences. These transgenes were all placed downstream of the upstream activating sequence (UAS) to achieve tissue specific gene expression. We first compared the relative microautophagy activities in various fly tissues using a Gal4 line driven by the heat shock promoter. Measuring the ratio between acidic (red) and neutral (green) Keima-CSPα signal showed high lysosomal translocation in intestine epithelial cells followed by ventral nerve chord, and the brain, while fat body had the lowest microautophagy activity (Fig. 6a, Extended Data Fig. 7a). The microautophagy activity of CSPα was modest in larval photoreceptor cells, but it could be enhanced when dCHIP was co-expressed (Fig. 6b). Thus, the CHIP’s function in microautophagy is conserved in flies.

**Fig. 6:**
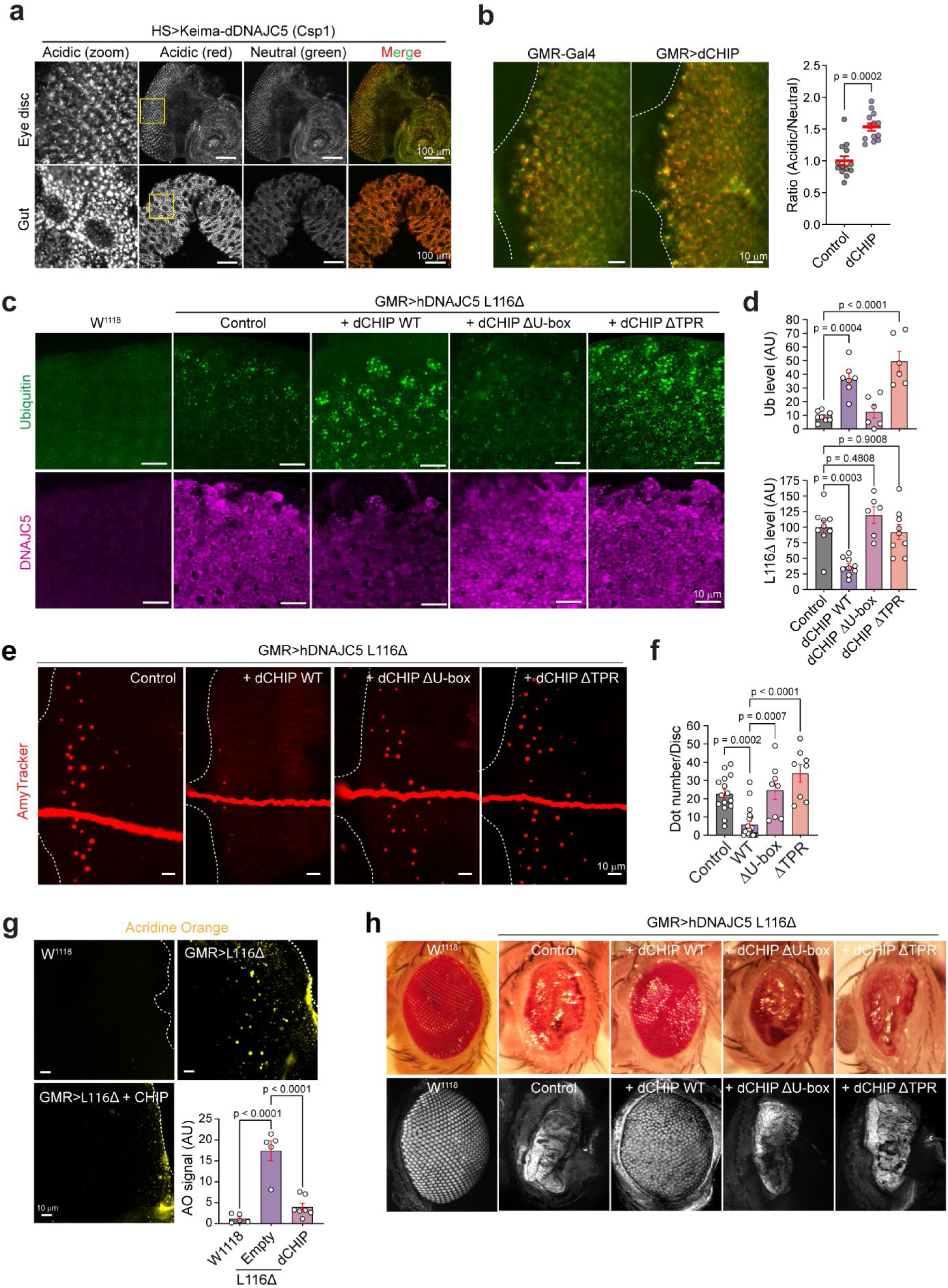
CHIP downregulates L116Δ and rescues lipofusin accumulation and neurodegeneration in a *Drosophila* CLN4 disease model. **a,** Representative confocal images of an eye disc and a segment of mid-gut from third insta larva of flies expressing Keima-dDNAJC5 (*Csp1*) by heat shock (HS)-Gal4. Scale bars, 100 µm. **b,** Confocal images of eye discs from third instar larva of GMR>Keima-dDNAJC5 (Control) or GMR>Keima-dDNAJC5; dCHIP (dCHIP). Scale bars, 10 µm. The graph shows the quantification of the acidic/neutral Keima signal ratio. Error bars represent means ± s.e.m., N=12 discs. P-value was determined by unpaired student’s t-test. **c,** Representative eye discs from larvae of the indicated genotypes were stained with anti-Ubiquitin (top panels) or DNAJC5 antibodies. Shown are maximum projected view of confocal sections of entire tissues. Scale bars, 10 µm. **d,** Quantification of the ubiquitin signal (top) or L116Δ levels (bottom) in individual eye discs. Error bars represent means ± s.e.m. P-values were determined by one-way ANOVA. **e,** Eye discs from larvae of GMR> Keima-DNAJC5 plus the indicated transgenes were stained with AmyTracker (2 µg/mL) and imaged. Scale bars, 10 µm. **f,** Quantification of the AmyTracker-positive punctae in e. Error bars represent means ± s.e.m. P-values were determined by one-way ANOVA. **g**, Eye discs from third insta larvae of the indicated genotypes were stained with acridine orange (1 µg/mL) and imaged. Scale bars, 10 µm. The graph shows the quantification of the experiment. Error bars represent means ± s.e.m. P-values were determined by one-way ANOVA. **h.** Eyes of flies with the indicated genotypes were either directly imaged by a CCD camera mounted on a dissecting microscope (top panels) or first fixed and then scanned by a confocal microscope (bottom panels). Confocal images in this figure were processed using maximum intensity projection to reconstruct the in-depth view of tissues.

We next crossed the UAS-CHIP lines to flies expressing human CLN4 L116Δ in larval photoreceptor cells via the GMR promoter. High level expression of human CLN4 L116Δ caused lipofuscin accumulation, enlarged lysosomes decorated with ubiquitin, and neuronal cell death ^27^, recapitulating phenotypes seen in CLN4 patient ^5^. Indeed, when imaginal eye discs from GMR>L116Δ third instar larva were stained with ubiquitin and DNAJC5 antibodies, photoreceptor cells expressing CLN4-L116Δ had more ubiquitin-positive puncta than those in WT flies (W*^1118^*) (Fig. 6c, d), and like in mammalian cells, many ubiquitin positive puncta were also positive for L116Δ (Extended data Fig. 7b). Confocal microscopy also detected many autofluorescent puncta resembling ceroid or lipofuscin. Interestingly, these structures could be stained by an amyloid specific dye, suggesting that they contain protein aggregates (Fig. 6e, Extended data Fig. 7c). As expected, acridine orange staining showed high level of cell death in L116Δ-positive but not in WT eye discs (Fig. 6g).

**Fig. 7:**
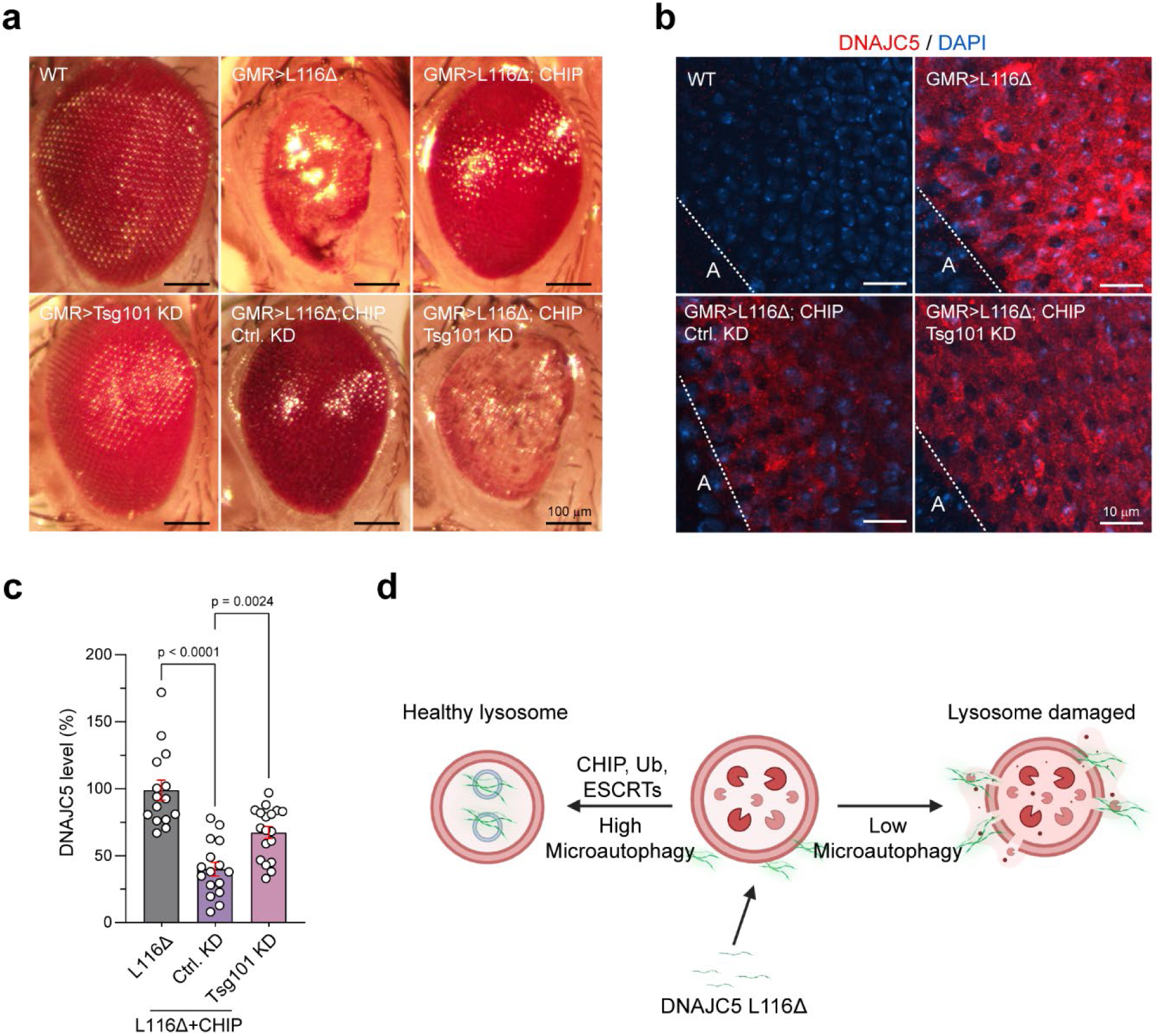
CHIP’s neuroprotective activity depends on ESCRT. **a**, Representative images of the eyes of flies with the indicated genotypes. Ctrl, Control; KD, knockdown. Scale bars, 100 µm. **b,** Eye discs from third instar larvae of the indicated genotypes were stained with DNAJC5 antibodies and DAPI (blue) to label DNA. Z-stack images were processed using maximum intensity projection. The white-dotted lines indicate the morphogenic furrow. A, anterior part. Scale bars, 10 µm. **c,** Quantification of the relative DNAJC5 fluorescence intensity in individual eye discs as shown in b. Error bars represent means ± s.e.m. P-values were determined by one-way ANOVA. **d,** A model depicts the role of CHIP- and ubiquitin-mediated microautophagy in counteracting CLN4 mutant-induced lysosome damage.

Co-expression of dCHIP in photoreceptor cells significantly reduced L116Δ protein level, while increasing lysosome-associated ubiquitin puncta (Fig. 6c, d), suggesting that dCHIP promotes lysosome-associated ubiquitination to down-regulate L116Δ. Neither dCHIP ΔU-box or dCHIP ΔTPR mutant could downregulate L116Δ, although only dCHIP ΔU-box failed to stimulate L116Δ-associated ubiquitination (Fig. 6c, d). Thus, additional factor(s) acting through the CHIP TPR domain are also involved in L116Δ downregulation. As expected, dCHIP but not the ΔTRP or ΔU-box CHIP mutant rescued lipofuscin accumulation (Fig. 6e, f) and mitigated L116Δ-associated neurodegeneration (Fig. 6g).

As expected from CHIP’s ability to downregulate L116Δ and reduce lipofuscin accumulation and cell death, WT dCHIP but not the U-box or TRP-deleted CHIP mutant suppressed the L116Δ-induced rough eye phenotype (Fig. 6h, Extended Data Fig. 7d). The rescue effect was comparable to that caused by HSC70 knockdown ^27^, which presumably prevents iron-sulfur cluster-mediated L116Δ aggregation (Extended Data Fig. 7d) ^24^. Conversely, knockdown of CHIP or the microautophagy regulator Tsg101 exacerbated the rough eye phenotype in L116Δ-expressing flies (Extended Data Fig. 7e). Notably, CHIP overexpression did not affect the rough eye phenotype induced by polyQ aggregates (Extended Data Fig. 7f), suggesting that its activity is specific towards lysotoxic proteins. Collectively, these data demonstrates that dCHIP enhances L116Δ-associated ubiquitination to downregulate L116Δ, and therefore, rescues lipofuscin accumulation and neurodegeneration in flies.

Lastly, we tested whether Tsg101-dependent microautophagy is required for CHIP’s lysoprotective activity. To this end, we generated GMR> hDNAJC5 L116Δ; dCHIP flies with or without Tsg101 shRNA. Knockdown of Tsg101 significantly reversed the eye improvement by dCHIP, although on its own, Tsg101 knockdown did not change the eye morphology (Fig. 7a). Immunostaining showed that dCHIP-mediated down-regulation of L116Δ was partially abolished when Tsg101 was knocked down (Fig. 7b, c). Together, these results suggested that dCHIP downregulates L116Δ at least in part via Tsg101-mediated microautophagy, which mitigates L116Δ-associated neurodegeneration.

## Discussion

LSDs are increasingly recognized as neurodegenerative disorders with lysosome deficiency as a common pathological hallmark. Accordingly, most CLN diseases are associated with recessive mutations in genes encoding lysosomal enzymes or factors essential for transporting these enzymes to lysosomes ^4^. In this regard, CLN4 is unique as it is linked to autosomal dominant mutations in *DNAJC5* presumed to confer a gain-of-toxic function, although the underlying mechanism has been unclear.

In i^3^Neurons bearing the L115R or L116Δ mutation in *DNAJC5*, we observed a dose-dependent lysotoxicity from the encoded mutant proteins, causing characteristic lysosomal abnormalities including enlarged lysosomes with damaged or ruptured membranes, diminished hydrolase activity, and AFSM accumulation. Our *in vitro* studies show that CLN4 mutants can oligomerize in an ATP- and cytosol-dependent manner, consistent with the reported role of cytosolic iron-sulfur clusters in CLN4 aggregation ^24^. Intriguingly, CLN4 aggregates generated *in vitro* can directly damage lysosomes from the cytosolic side in permeabilized cells. Since several studies reported that endocytosed protein aggregates can also damage lysosomal membranes but from the luminal side ^42, 43^, lysosome membranes may be intrinsically vulnerable to protein aggregate-induced membrane instability.

Notably, it was reported that polyQ-containing protein aggregates can disrupt and deform the endoplasmic reticulum membranes ^44^. How cytosolic protein aggregates compromise distinct membrane compartments remains to be elucidated, but these processes likely involve specific membrane adaptors that bring protein aggregates to the target membrane. Once in proximity to the membranes, protein aggregates may destabilize the lipid bilayer directly, form a membrane-embedded pore through oligomerization, or damage membranes via reactive oxygen species (ROS) generation. Lysosomal membrane rupture is known to trigger cascading cellular damage, activating inflammation, and releasing lysosomal enzymes, which ultimately leads to cell death ^45, 46^.

Cells deploy several protective mechanisms to maintain lysosomal integrity, which include ESCRT-mediated membrane repair and lysophagy ^36, 47^. Our findings identify ubiquitin-mediated microautophagy as an additional organelle damage control mechanism that safeguards lysosome homeostasis (Fig. 7d). Microautophagy can target toxic cytosolic protein aggregates for degradation, therefore, protects lysosomes from aggregate-induced membrane damage. Additionally, microautophagy may facilitate membrane repair, since ESCRT-mediated MVB formation is known to remodel lyososomal membranes, which can aid in the repair of damaged lysosomal membranes ^32–35^.

Our findings indicate that CHIP-dependent ubiquitination at CLN4-positive lysosomes serves as a molecular linchpin in ubiquitin-dependent lysosomal protection. In non-neuronal cells, CHIP-mediated microautophagy directs aggregates and possibly damaged membranes toward the degradation pathway, which explains the lack of significant lysosomal defect in fibroblast cells bearing CLN4 disease mutations ^30^. However, during neuronal differentiation, this protective function appears impaired when CLN4 mutants are present, as CHIP becomes sequestered within insoluble aggregates, which correlates with increased CLN4 aggregation and lysosome destabilization during i^3^Neurons differentiation. Whether CHIP activity is similarly downregulated during neuronal differentiation in animals is unclear. However, since CHIP expression is relatively low in both excitatory and inhibitory neurons according to the Human Protein Atlas (proteinatlas.org) ^48^, we presumed that diminished CHIP expression/function in neurons may cause the vulnerability to CLN4-induced lysotoxicity. Consistent with this notion, ectopic CHIP expression can partially restore lysosomal integrity and cell viability in CLN4 i^3^Neurons and mitigate CLN4 pathology in a *Drosophila* model of CLN4 disease. The rescue effect of CHIP in *Drosophila* is better than in i^3^Neurons, probably because CHIP was co-expressed with L116Δ in flies using the same promoter, while in i^3^Neurons, CHIP expression driven by the *Synapsin* promoter lags behind L116Δ aggregation.

CHIP has long been recognized as a critical component of the cellular protein quality control system, since it can interact with cytosolic chaperones HSC70 and HSP90. Consequently, CHIP can preferentially ubiquitinate misfolded or aggregation-prone proteins and target them for degradation ^49^. CHIP’s vital role in quality control and homeostasis regulation is underscored by numerous genetic links to neurodegenerative diseases such as spinocerebellar ataxia ^49^. Intriguingly, recent studies suggested that CHIP’s activity is regulated by a monomer-to-dimer switch: dimeric CHIP mediates chaperone-assisted turnover via the proteasome, while monomeric CHIP promotes the turnover of membrane-bound proteins, as observed in *C. elegans* ^50^. Whether CHIP’s role in lysosome homeostasis regulation requires the monomeric or dimeric form and how CHIP-associated chaperones aid in this process remain to be tested.

The relevance of our findings to broader neurodegenerative disease mechanisms is compelling. Lysosomal membrane damage is commonly reported in neurodegenerative diseases such as Parkinson’s and Alzheimer’s diseases, where internalized protein aggregates are major contributors to membrane destabilization and lysosome leakage ^42, 43, 51^. These observations suggest that CHIP’s role in microautophagy may also exploited to develop therapeutics for lysosome damage-associated disorders. Supporting this idea, CHIP overexpression was shown to promote the degradation of an aggregated form of α-synuclein in human H4 neuroglioma cells ^52^, though whether this process involves microautophagy is unclear.

In summary, our study provides novel insights into the pathological role of CLN4-associated mutant DNAJC5 aggregates, establishing lysosomal damage as a central feature of the disease. Our work reveals a critical role for CHIP-dependent ubiquitination and microautophagy in lysosomal damage control and suggests that modulation of this lysosome protection mechanism may hold therapeutic promise for ameliorating lysosomal dysfunction in CLN4 and other aggregate-associated neurodegenerative diseases.

## Materials and Methods

### Cell lines, plasmids, antibodies, and other reagents

U-2 OS (U2OS) and 293T cells were purchased from ATCC (catalog no. HTB-96 and CRL-3216). Cells were grown in DMEM (Corning, catalog no. 10-013-CV) supplemented with 10% FBS (Corning, catalog no. 35-011-CV) and 100 µg/mL penicillin-streptomycin (Gibco, catalog no. 15140-122), and maintained at 37 °C, 5% CO_2_, and 95% humidity. Lipofectamine 2000 (Invitrogen, catalog no. 11668027) and TransIT-293 Transfection Reagent (Mirus Bio, catalog no. MIR 2700) were used for transfection in U2OS and 293T, respectively, according to the manufacturer’s protocol. For siRNA transfection, Lipofectamine RNAiMAX (Invitrogen, catalog no. 13778075) was used according to the manufacturer’s protocol. All plasmids used in this study were constructed by standard molecular biology methods and sequenced confirmed. All plasmids will be deposited to Addgene. Plasmids, antibodies and reagents used in this study are listed in Supplementary Table 2.

### Lentivirus production

To prepare lentiviruses, HEK293FT cells (Thermo Fisher Scientific, catalog no. R70007) were transfected with pVSV-G (Addgene #8454), psPAX2 (Addgene #12260), and lentiviral expression vectors (e.g., pLenti CMV Hygro or FSW) in a 1:1.5:2 mass ratio. After 24 h, the media were replaced with fresh media and incubated for 48 h to collect viruses. The viral supernatants were harvested, centrifuged at 300 g for 10 minutes, filtered through a 0.45 µm PVDF syringe filter unit (Sigma-Aldrich, catalog no. SLHVM33RS), and concentrated ∼100x using Lentivirus Precipitation Solution (ALSTEM, catalog no. VC100) according to the manufacturer’s protocol. The resulting virus pellets were resuspended in DPBS (Gibco, catalog no. 14190-144) and Virus Protection Medium (ALSTEM, catalog no. VF050), aliquoted, flash-frozen in liquid nitrogen, and stored at −80 °C.

### CRISPR-CAS9 gene editing

*CHIP* and *CHMP6* knock-out in 293T and *CHIP* knock-out in U2OS cell lines were carried out by infection with lentiviruses carrying Cas9 and the corresponding sgRNA in pLenti CRISPRv2 (Addgene #52961), as described ^39^. The following gRNAs, designed using the ATUM website (https://www.atum.bio/eCommerce/cas9/input), were used: sg-hCHIP forward: 5’-caccgTGTATTACACCAACCGGGCC-3’, sg-hCHIP reverse: 5’-aaacGGCCCGGTTGGTGTAATACAc-3’; sg-hCHMP6 forward: 5’-caccgGCTCAAGAAGAAGCGATACC-3’, sg-hCHMP6 reverse: 5’-aaacGGTATCGCTTCTTCTTGAGCc-3’. Infected cells were cultured for three days followed by puromycin (gibco, catalog no. A1113803) selection (0.4 µg/mL for 293T and 1 µg/mL for U2OS cells) until the parallel non-infected control cells were all killed. After one passage in medium without puromycin, cells were infinitely diluted and seeded at 1 cell per well into 96 well plates either manually or using a BD Fusion cell sorter. Knock-out clones were verified by immunoblotting.

To generate L115R and L116Δ knockin iPSC cells, 0.8 million HT727D iPSCs ^28^ were transfected with premixed 60 pmol HiFiCas9 V3 (IDT, catalog no. 1081061), 60 pmol dCas9 V3 (IDT, catalog no. 1081067), 200 pmol sgRNA (Synthego), and 200 pmol single strand oligodeoxynucleotides (ssODN) (IDT) using Nucleofector 4D with buffer P3 and program CA-137 (Lonza, catalog no. V4XP-3024) and then plated onto one well of rhLaminin-521 (gibco, catalog no. A29249) coated 6-well plate with StemFlex (gibco, catalog no. A33493-01), RevitaCell (gibco, catalog no. A2644501) and HDR enhancer V2 (IDT, catalog no. 10007921). Transfected cells were cultured in 32 °C incubator for 3 days before moving to 37 °C incubator, and fresh medium was changed daily after transfection. Single-cell subcloning was done using manual serial dilution to 1 cell/96-well Matrigel coated plate with StemFlex and CloneR2 (Stem Cell Technologies, catalog no. 100-0691). 10 days later, single-cell clones were picked and confirmed by genomic PCR, Sanger sequencing, and ICE analysis (Synthego). Besides L115R and L116Δ mutation knock-in clones, biallelic DNAJC5 knockout clones and unedited isogenic control clones were also saved. The sgRNA target, single-stranded oligodeoxynucleotides (ssODNs), and PCR primers used for p.L115R (c.344T>G) and p.L116Δ (c.346-348Δ) mutation knock-in in DNAJC5 are listed below: p.L115R (c.344T>G), sgRNA target: AGCAGTAGCAGCACGTGAGG, ssODN: aacaccaccttcttctccccccagGCCCTGTTTGTCTTCTGCGGCC**G**CCTGACGTGCTGCTACTGCTGCTGCT GTCTGTGCTGCTGCTTCAACTGCTGCT (Note: this sequence contains a L116 CTC to CTG silent mutation to avoid recutting after L115R KI).

p.L116del (c.346-348del), sgRNA target: AGCAGTAGCAGCACGTGAGG, ssODN: aacaccaccttcttctccccccagGCCCTGTTTGTCTTCTGCGGCCTGACGTGCTGCTACTGCTGCTGCTGTCTGTGCTGCTGCTTCAACTGCTGCT (contains L115 CTC>CTG silent mutation)

PCR forward primer: TGCTTTTCTTTAAGCTGCGGG

PCR reverse primer: TACCTCAGGGTCCACGTTCA

### Cell lysis, membrane fractionation, and immunoblotting

To lyse cells, ∼2-3 million 293T cells were washed with phosphate-buffered saline (PBS) two times and then treated with 300 µL the NP40 lysis buffer containing 0.5% Nonidet P-40 (Sigma-Aldrich, catalog no. 56741-50ML-F], 50 mM Tris-HCl, pH 7.4, 150 mM NaCl, 2 mM MgCl_2_, 1 mM EDTA, 1 mM TCEP (Thermo Fisher Scientific, catalog no. 20490), and a protease inhibitor cocktail. Cells were incubated on ice for 15 min with occasional mixing. The lysates were cleared by centrifugation (17,000 g, 10 min at 4 °C). The supernatant (the NP40-soluble fraction) was mixed with 4x Laemmli buffer (BioRad, catalog no. 1610747) and then heated at 95 °C for 5 min. The remaining pellets were washed once with the NP40 lysis buffer and then centrifuged at 17,000 g for 10min at 4 °C. The pellets were resuspended in 100 µL PBS by gentle pipetting and then mixed with equal volume of pre-heated 2x Laemmli buffer (BioRad). The samples were immediately heated at 95 °C for 20 min to obtain the NP40-insoluble fraction.

Cytosol-membrane fractionation was performed as described previously ^22^. Cells harvested in PBS were pelleted by centrifugation at 500 g for 5 min at 4°C. Cell pellets were treated with a permeabilization buffer (PB) containing 0.025% digitonin (Sigma-Aldrich, catalog no. D141), 230 mM potassium acetate (Sigma-Aldrich, catalog no.P1190), 10 mM sodium acetate (Sigma-Aldrich, catalog no. S2889), 50 mM HEPES, pH7.3, 5 mM MgCl_2_, 1 mM EGTA (Sigma-Aldrich, catalog no. 324626), 1 mM TCEP (Thermo Fisher Scientific), and a protease inhibitor cocktail on ice for 5 min. Plasma membrane permeabilization was confirmed by trypan blue (Thermo Fisher Scientific, catalog no. 15250061) staining. Cells were spun at 17,000 g for 5 min. The supernatant was saved as the cytosol fraction. The resulting membrane pellets were washed with 1x PB buffer followed by centrifugation and then further lysed by a CHAPS lysis buffer (1% CHAPS [Sigma-Aldrich, catalog no. 10810118001], 50 mM HEPES, pH 7.4, 100 mM NaCl, 1 mM TCEP, and protease inhibitors). The lysates were cleared by centrifugation at 17,000 g for 5 min and the supernatant fractions were collected as membrane fraction.

For immunoblotting, samples were loaded onto 4-12% Bis-Tris gels (Invitrogen) followed by SDS-PAGE with MES SDS Running Buffer (Invitrogen, catalog no. B0002). The proteins were transferred to Nitrocellular membranes (0.45 µm, BioRad, catalog no. 1620115), which were stained using the Ponceau S. solution (Sigma-Aldrich, catalog no. P7170). Membranes were then blocked with PBS containing 5% non-fat milk, washed with PBS three times, and then incubated with primary antibodies in 5 % BSA (Sigma-Aldrich, catalog no. A9418) in PBS supplemented with sodium azide (0.03%) for overnight in 4 °C. The membranes were washed three times with PBS and further incubated either with goat anti-Mouse DyLight™ 680, goat anti-Rabbit DyLight™ 800 (Thermo Fisher Scientific), or HRP-labeled goat anti-Mouse or Rabbit IgG (Sigma-Aldrich) at room temperature for 1 h. After thorough wash with PBS for three times, fluorescence or chemiluminescence signal was detected using a ChemiDoc MP scanner (Biorad) and quantified using the ImageLab software (BioRad).

### Immunoprecipitation

For co-immunoprecipitation assay, 293T cells transfected with FLAG-DNAJC5 were fractionated into cytosol and membrane fractions as described above. ANTI-FLAG® M2 Affinity Gel (Sigma Aldrich, catalog no. A2220) was added to the fractions and incubated for 1 h at 4 °C. A portion of the extracts were saved as input before adding the beads. The beads were washed by PBS three times and the proteins bound were eluted by 1x Laemmli buffer and heating at 95 °C for 5 min before SDS-PAGE and immunoblotting analysis.

Denatured immunoprecipitation was performed to detect ubiquitination on DNAJC5. To this end, the cells were transfected or infected with lentiviruses to achieve the expression of tagged-DNAJC5 variants. For lentivirus-infected cells, drug selection was conducted to make cells stably expressing these DNAJC5 variants. Cells were lysed in a buffer (150 µL) containing 0.5 % NP40, Tris-HCl 7.4 50 mM, 150 mM NaCl, 2 mM MgCl_2_, 1 mM DTT, 2 mM NEM and protease inhibitor. After centrifugation, cleared cell extracts were adjusted with SDS and DTT to contain 1% SDS and 5 mM DTT. Cell lysates were heated at 95 °C for 5 min. Lysate were diluted with the NP40 lysis buffer by 10-fold and then centrifuged. The cleared lysates were incubated with FLAG beads or DNAJC5 antibody-containing beads to purify DNAJC5. Bound proteins were washed and then eluted with 1x Laemmli buffer by heating at 95 °C for 5 min. Eluted proteins were analyzed by SDS-PAGE and immunoblotting with anti-ubiquitin.

### Human iPSC culture and neuronal differentiation

Human WT and CLN4 mutation-bearing iPSC cells were transduced with a doxycyclin-inducible NGN2 as described previously ^53^. These cells were cultured on Matrigel-coated dishes in StemFlex medium (Thermo Fisher Scientific) according to the manufacturer’s instruction. Briefly, cells were passaged when they reach 70% confluent using StemPro Accutase (gibco, catalog no. A11105-01) Cells were seeded into Matrigel (Corning, catalog no. CLS354277)-coated dishes with a density of 10,000 cells per cm^2^ in the presence of 1 µM Chroman 1 (MedChemExpress, catalog no. HY-15392). The chroman 1 was removed on the following day and medium change was performed every day. If necessary, EZ-Lift Stem Cell Passaging Reagent (Sigma-Aldrich, catalog no. SCM139) was used to eliminate spontaneously differentiated cells from the culture according to the manufacturer’s protocol. Differentiation of iPSCs to i^3^Neurons was performed as previously described ^53^ with minor modifications. On day 0, iPSCs were seeded to Matrigel-coated dishes at 80,000 cells per cm^2^ using the KnockOut DMEM/F-12 (Thermo Fisher Scientific, catalog no. 12660012)-based induction medium containing 1x N-2 supplement (Thermo Fisher Scientific, catalog no. 17502048), 1x GlutaMax (gibco, catalog no. 35050061), 1x non-essential amino acids (NEAAs; gibco, catalog no. 11140050), 1 µM Chroman 1 and 2 µg/mL doxycycline (Sigma-Aldrich, catalog no. D5207). Medium change was performed on Day 1, 2 and 3 using induction medium without Chroman1. On day 3, 1 µM 5-Fluore-2’-deoxyuridine (Sigma-Aldrich, catalog no. F0503) and 1 µM Uridine (Sigma-Aldrich, catalog no. U3003) were added for overnight treatment to eliminate undifferentiated cells. On day 3, ibidi 8-well glass chamber (for imaging) or corning dishes (for biochemical assays) were coated with 100 µg/mL Poly-L-Ornithine (Sigma-Aldrich, catalog no. P3655) for overnight at 37 °C. Next day (day 4), the wells were washed with DPBS twice and further coated with 1 µg/mL of rhLaminin-521 (gibco) for 1 h at 37 °C. At the day 4, the differentiated neurons were dissociated using Accutase, counted and plated with a density of 100,000 cells per cm^2^ (for imaging) or 200,000 cells per cm^2^ (for biochemical assays) using neuronal culture medium containing 1:1 mix of BrainPhys medium (STEMCELL, catalog no. 5790) and KnockOut DMEM/F-12 supplemented with 1x N21-Max (R&D Systems, catalog no. AR008), 10 ng/mL BDNF (Peprotech, catalog no. 450-02), 10 ng/mL GDNF (Peprotech, catalog no. 450-10), 10 ng/mL NT-3 (Peprotech, catalog no. 450-03), 0.5 µg/mL rhLaminin (gibco) and 2 µg/mL doxycycline (Sigma-Aldrich). At day 6, the medium was changed with fresh neuronal culture medium without KnockOut DMEM/F-12 and Doxycyclin. Since then, half of the medium was removed every 3 days, and an equal volume of fresh medium was added for maintenance. If necessary, iPSC or day 4 i^3^Neurons were frozen using CryoStor® CS10 medium (STEMCELL, catalog no. 100-1061) according to the manufacturer’s protocol and kept in liquid nitrogen tank for storage.

### Transmission Electron Microscopy

Cells were fixed in 2.5 % Glutaraldehyde, 2 % Formaldehyde, in 0.1 M Cacodylate/2 mM Calcium chloride pH 7.4 (CacCl) for 15 min at room temperature followed by 45 min incubation on ice. Coverslips were washed 3 times for 5 min each with CacCl, post-fixed for 30 min in 0.5 % Osmium tetroxide/0.5% potassium ferrocyanide in the same buffer, washed and treated with 1 % tannic acid for 30 min. Cells were washed 2 times with CacCl, 2 times with 50 mM sodium acetate (pH 5.2) and stained overnight with 2% Uranyl acetate. After washing 2 times with the acetate buffer and 2 times with water, the samples were dehydrated through a series of increasing concentration of ethanol (50%, 75%, 90%, 3 times in 100% anhydrous) and embedded in EMBed_812_ epoxy resin (EMS, catalog no. 14120)). After resin polymerization, the coverslip was removed by hydrofluoric acid. Sample blocks were cut out and mounted on a holder. Ultrathin sections (80 nm thick) were cut parallel to the plane of the coverslip and mounted on formvar/carbon coated EM grids. Sections were stained with lead citrate and imaged in FEI Tecnai 20 transmission electron microscope operated at 120 kV. Images were recorded on AMT Nanosprint 12 wide field CCD camera.

### Cell staining, confocal imaging, and data analysis

mKeima fluorescence and autofluorescence (AFSM) were detected in live cells following previously reported protocols ^22^. For immunostaining, i^3^Neurons or U2OS cells cultured on µ-Slide 8 Well high Glass Bottom (ibidi, catalog no. 80807) were fixed in PBS containing 4 % paraformaldehyde (Thermo Fisher Scientific, catalog no. 28908) for 10 min at room temperature. Cells were then washed with PBS twice and permeabilized with a PBS-based staining solution containing 0.2 % saponin (Sigma-Aldrich, catalog no. 47036) and 10 % FBS for 10 min at room temperature. For methanol fixation, cold methanol was added to PBS-washed cells. Cells were incubated at −20 °C for 10 min and washed with PBS three times and blocked with 5% FBS in PBS for 10 min at room temperature. Cells were stained by primary antibodies diluted in the staining solution overnight at 4 °C and washed with PBS three times. Alexa Fluor^®^ labeled secondary antibodies (Thermo Fisher Scientific) were diluted in the staining solution and added to cells for 1 h at room temperature. Where indicated, 1 μg/mL DAPI (Sigma-Aldrich, catalog no. D9542) was included in the staining solution to label nuclei. The slides were washed three times with PBS. The stained slides were imaged by either LSM780 laser scanning confocal microscopy (Zeiss) or CSU-W1 SoRa spinning disk super-resolution microscopy (Nikon).

For LysoTracker and Magic Red staining in live i^3^Neuron, LysoTracker™ Red DND-99 (1: 5,000; Invitrogen, catalog no. L7528) or Magic Red (1: 250; Antibodies Inc., catalog no. 938) was added to cells and stained for 30min. For the DQ-BSA assay, i^3^Neurons were incubated with DQ red BSA (Invitrogen, catalog no. D12051) at 40 µg/mL for 6 h. Where indicated, Hoechst33442 (1:10,000; Invitrogen, H3570) was added to the media for labeling nuclei. Random-fielded images were collected as z-section to cover entire cell volume using the Nikon CSU-W1 SoRa confocal microscope equipped with 60x TIRF objective (NA=1.6) and a heating and CO_2_-perfused chamber.

For data quantifications, ImageJ software was used to merge all the z-sections, generating maximum projected images. Fluorescence intensity was measured by using ROIs covering the soma of i^3^Neurons. A portion of soma with no signal in each image was selected for background subtract. The total adjusted intensity was calculated by multiply individual mean intensity with the cell area. To measure the lysosome volume or the area of immunostained signals, images were subject to programed image thresholding. Automate single particle analysis was used to identify lysosomes and measure their size.

For cell viability assay in i^3^Neuron, the LIVE/DEAD™ Viability/Cytotoxicity Kit (Invitrogen, catalog no. R37601) was used according to the manufacturer’s protocol. The number of live neurons (green) and dead neurons in randomly selected fields were determined by using a threshold and watershed function in ImageJ. The cell death index (normalized red) was calculated as the number of red cells divided by the number of green plus red cells (Total cells). For 3-D visualization, we used Imaris (Oxford instruments). Some images were processed using Nikon NIS-element plugin Denoise.ai for background reduction.

For plate reader-based fluorescence measurement, i^3^Neurons cultured in 96 well plates were measured by GloMax® Explorer Multimode Microplate Reader (Promega). Wells containing no staining dyes were used for background subtraction. For area or intensity analyses, untransfected cells next to transfected cells were used as internal controls to measure the relative fold-change.

### Membrane fractionation and lysosome isolation

Lysosome Enrichment Kit (Thermo Fisher Scientific, catalog no. 89839) was used for fractionating organelles from i^3^Neurons according to the manufacturer’s protocol. Briefly, two p100 dishes containing i^3^Neurons at day16 per condition were washed with DPBS twice, and neurons were harvested by centrifugation at 1,000 g for 10 min. After removal of DPBS, cells were resuspended in 500 µL Buffer A with protease inhibitors, vortexed for 5 seconds, and incubated on ice for 5 min. Plasma membranes were broken by 60 strokes using a 2 mL glass Dounce homogenizer with a tight pestle. 500 µL buffer B with protease inhibitors was added. The extracts were cleared by centrifugation at 500 g for 10 min at 4 °C. The resulting supernatants with 15% of OptiPrep™ cell separation media were divided into three equal portions and each portion was overlayed on a discontinued OptiPrep™ media gradient (17, 20, 23, 27, and 30 %) in ultra-clear centrifuge tubes (Beckman Coulter, catalog no. 344062) following the provided protocol. After ultracentrifugation at 145,000 g (37,600 rpm) for 2 h at 4 °C, 0.5 mL fractions were carefully collected from the top of gradients and gently mixed with 1 mL PBS. The fractions were centrifuged at 18,000 g for 30 min at 4 °C. The supernatant fractions were removed. PBS (0.2 mL) was added to each fraction to resuspend the membrane pellet. The same fractions were combined and then centrifuged at 18,000 g for 30 min at 4 °C again. The supernatant fractions are removed and stored at −80°C until immunoblotting and mass spectrometry analyses.

To immune purify lysosomes, lyso-IP method was used as previously described with modifications ^54^. Briefly, i^3^Neurons at day6 were infected by lentivirus to express TMEM192-HA with CHIP-Myc or empty vector. At day16, neurons were washed once with Tris-buffered saline (TBS) and harvested by centrifugation at 450 g for 5 min. TBS is removed and cell pellets were resuspended with cold fresh TBS with protease inhibitors and lysed by 20 strokes using a 2 mL glass Dounce homogenizer with a tight pestle. The lysates were transferred and mixed with 8 mM CaCl_2_ and vortexed. After centrifugation at 1,150 g for 3 min in 4 °C, supernatants were carefully transferred into new tubes and used for immunoprecipitation using Anti-HA Magnetic Beads (Thermo Fisher Scientific, catalog no. 88836) pre-equilibrated with the same buffer. A small fraction was saved before adding the beads for input. The beads were incubated on a rocker for 4 h in 4 °C and washed with TBS with 8 mM CaCl_2_ and protease inhibitors six times using DynaMag™-2 Magnet (Invitrogen, catalog no. 12321D). Lysosomes were eluted and lysed by heating at 65 °C for 10 min in the presence of 1x Laemmli buffer.

### Organelle proteomics sample preparation

Membrane pellets prepared as described above were reconstituted in ice-cold lysis buffer (500 mM NaCl, 0.1% SDS, 1% Triton, 5 mM TCEP) and sonicated using QSonica sonicator (QSonica, catalog no. Q800R) for 10 min in an ice-cold water bath with alternating 40 s-on and 20 s-off cycles. Protein concentration was determined using a Bio-Rad Detergent Compatible (DC) protein assay. Protein reduction, alkylation, and digestion were conducted with an automated SP3 method in an automated KingFisher sample preparation system as described previously ^55^. Briefly, samples were reduced with 5 mM TCEP at room temperature for 40 min, alkylated with 15 mM IAA for 40 min at 37 °C, and quenched with 15 mM DTT for 15 min at 37 °C. Acetonitrile (ACN) was added to 80% percentage in volume in each sample, and 10 µL of SP3 beads (Cytiva) was added in the sample followed by 10 min of incubation to induce protein binding. Beads were washed three times with 95% ACN, two times with 70% ethanol, and then released in 100 µL of 50 mM ammonium bicarbonate buffer. Proteins were digested with Trypsin/Lys-C mix (1:25, enzyme: protein) for 1 h at 47 °C. Beads were washed again in 100 µL of LC-MS grade water. Both supernatants of peptides elution were combined. Residual detergents from digested peptides were tested and removed using the ContamSPOT assay prior to LC-MS analysis, as described previously ^56^. Peptide samples were dried down under SpeedVac and kept in −30 °C.

### LC-MS/MS analysis

Dried peptide samples were reconstituted in 0.1% formic acid (FA), 2% ACN in LC-MS grade water and analyzed on a Dionex Ultimate 3000 RSLCnano system coupled with a Thermo Fisher Scientific Scientific Q-Exactive HF-X Orbitrap mass spectrometer. Peptides were separated on an Easy-Spray PepMap RSLC C18 column (2 µm, 100 Å, 75 µm × 50 cm) with a 180 min LC gradient and a flow rate of 0.25 µL/min. Mobile phase A was 0.1% FA in water, and mobile phase B was 0.1% FA in ACN. Samples were analyzed in a staggered data-independent acquisition mode with 75 sequential scans and an *m/z* 8.0 isolation window. MS1 scanned from *m/z* 400 to 1,000 with a resolving power of 60K, an automatic gain control (AGC) target of 1E6, and a maximum injection time (maxIT) of 60 ms. MS2 resolving power was 15K, AGC target was 2E5, and maxIT was 40 ms. Normalized collision energy was 30%.

### Proteomics data analysis

Proteomics raw data was analyzed in the Spectronaut software (v18.1, Biognosys). Proteins and peptides were identified with a 1% false discovery rate cut off using the Swiss-Prot Homo sapiens database and neuron-specific contaminant library ^57^. Missed cleavages were set up to 2 and variable modifications up to 3. Cysteine carbamidomethylation was set as fixed modifications, and methionine oxidation, and protein N-terminal acetylation was set as variable modifications. Precursor intensities below 1000 were removed from the Spectronaut protein report file. Statistical analysis was conducted using a two-tailed Student’s t-test.

### In vitro lysosome-damaging assay

To test the lysosome damaging activity of CLN4 mutants, 25,000 cells of U2OS cells were seeded in an 8-well Ibidi cell chamber two days before the experiment. We then labeled the cells with Lysotracker^TM^ Red DND-99 (Thermo Fisher Scientific) at 1 µM for 30 min at 37 °C. Cells were washed four times with ice-cold phosphate-buffered buffer saline containing 2 mM MgCl_2_ (PBS-Mg) and then treated with the same buffer (300 µL) containing 100 units of Streptolysin O (Sigma-Aldrich, catalog no. SAE0089) on ice for 10 min. Cells were then washed three times with the PBS-Mg buffer and the incubated with 200 µL reagent mixture containing 150 µL PBS-Mg, 50 µL cow liver cytosol or buffer, 2 mM ATP, 1 mM DTT, 0.5 µM LysoTracker Red, a cell impermeable dye and a protease inhibitor cocktail. WT or CLN4 mutant proteins were added at 2 µM. Cells were incubated at 37 °C for 30 min and then imaged by a Nikon CSU-W1 SoRa super-resolution confocal microscope.

To measure lysosome by Dextran leakage, U2OS cells were first loaded with Dextran Alexa Fluor™ 568 10,000 MW (100 µg/mL; Invitrogen, catalog no. D22912) at 37 °C for 4 h, and then incubated in a Dextran-free medium for 3 h. Cells were then labeled with a LysoTracker^TM^ Green dye (Invitrogen, catalog no. L7526) at 1 µM for 30 min before treated with Steptolysin O.

### Recombinant protein purification

To purify recombinant human DNAJC5 (hDNAJC5) proteins, hDNAJC5 WT, L115R and L116Δ proteins containing a TEV cleavage site between a GST tag and the protein were expressed in BL21(DE3) competent *E.coli* (NEB, catalog no. C2527) by adding 0.5 mM IPTG to 2 liters LB cultures at 16 °C for overnight. Cells were harvested by centrifugation at 6,000 g for 20 min at 4 °C, and the pellets were resuspended in 35 mL PBS containing 2 mM TCEP and protease inhibitors. Cells were lysed by sonication (30 % amplitude) on ice with the setting of 10 sec ON / 20 sec OFF for 10 min of total ON time. The lysates were cleared by centrifugation at 40,000 g (∼18,000 rpm) for 30 min in 4 °C and the supernatants were incubated with 2 mL (1 mL beads bed volume) of Glutathione Sepharose™ 4 Fast Flow (Cytiva, catalog no. 17513202) pre-equilibrated with PBS on a shaker for overnight. The beads were transferred to glass chromatography columns (Bio-Rad) and washed three times by 15 mL of PBS with 2 mM TCEP. To elute proteins from the beads, 1 mL PBS containing 400 unit of biotin-tagged TEV protease (Sigma-Aldrich, catalog no. SAE0118) was added to the beads and incubated at 4 °C overnight. Elutes containing proteins were collected by gravity flow and further incubated with 100 μL of 1:1 mixture of Glutathione Sepharose (Cytiva) and High-Capacity Streptavidin Agarose (Thermo Fisher Scientific, catalog no. 20357) for 1 h at 4 °C to remove the TEV protease. The protein purity was confirmed by SDS-PAGE and Coomassie staining. The proteins were aliquoted, flash-frozen in liquid nitrogen and stored at −80 °C.

### Genome-wide CRISPR/CAS9 knockout screen

The GeckoV2 library was purchased from Addgene (1-000-000-048) and amplified according to the online protocol from Feng Zhang’s lab ^39^. The complexity of the sgRNA library was verified by high-throughput sequencing by the NIDDK Genomic Core. Lentiviruses containing the sgRNA library and Cas9 were generated and used for transduction via spinfection. Briefly, 60 million of 293T stably expressing mKeima-tagged human DNAJC5 WT or ΔJ mutant supplemented with 8 μg/mL polybrene (Sigma-Aldrich, catalog no. TR-0003) were seeded in two 12-well plates at a density of 3 million cells per well. Concentrated GeCKO v2 lentivirus (library A) was added to each well at a multiplicity of infection (MOI) at 0.3. Cells were then spun at 1,000 g at room temperature for 2 h followed by incubation at 37 °C in a humidified incubator for 1 h. After the medium was removed, fresh growth medium was added, and cells were incubated for 48 h before the start of the selection for lentiviral integration using puromycin (0.3 μg/mL). The transduced cells were subcultured in medium supplemented with puromycin every 2 days for a total of 8 days and 80 million cells were maintained for each passage. After puromycin treatment, cells were recovered in a medium lacking puromycin for 24 h before cell sorting. In total, 100 million cells were sorted into mKeima neutral (3% of total cells) or mKeima acidic (97% of total cells) cell populations by a FACSAria™ Fusion Flow Cytometer (BD Bioscience). Genomic DNA was extracted from each cell population using a QIAGEN Blood Maxi kit (for mKeima acidic cells; Qiagen, catalog no. 51192) or QIAGEN Blood Midi kit (for mKeima neutral cells; Qiagen, catalog no. 51183) according to the manufacturer’s protocols. sgRNAs sequences were amplified from genomic DNA samples using the Herculase II Fusion DNA Polymerase (Agilent, catalog no. 600675) in two PCR steps as follows. In the first PCR, the genomic region containing sgRNAs were amplified from ∼250 μg and ∼20 μg of total DNA from acidic and neutral samples respectively, using the following primers: Forward: 5′-AATGGACTATCATATGCTTACCGTAACTTGAAAGTATTTCG-3′; Reverse: 5′-TTCAAAAAAGCACCGACTCGGTGCCACTTTTTCAAGTTGATAAC-3′. In total, 32 and three PCR reactions for acidic and neutral samples respectively were performed in parallel, with 8 μg genomic DNA in each reaction using Herculase II Fusion DNA Polymerase (Agilent) for 18 cycles and the resulting PCR products were combined. In the second step PCR, 5 μL of first PCR product was used in a 100 μL reaction volume and 12 PCR cycles were used. The primers used for the second PCR include stagger sequences of variable lengths and a 6 bp barcode for multiplexing of different biological samples. The following PCR primers were used in the second step: Forward: 5′-AATGATACGGCGACCACCGAGATCTACACTCTTTCCCTACACGACGCTCTTCCGATCT-(1-7bp variable length sequences)-(6bp barcode)-TTGTGGAAAGGACGAAACACCG −3’; Reverse: 5′-CAAGCAGAAGACGGCATACGAGAT-(6bp barcode)- GTGACTGGAGTTCAGACGTGTGCTCTTCCGATCTTCAAGTTGATAACGGACTAGCC-3′. The final PCR products were gel extracted, quantified, and sequenced using a NovaSeq sequencer (Illumina) by the NHLBI DNA Sequencing and Genomics Core. sgRNA sequences were obtained per sample by extracting 20 bps followed by the index sequence of “TTGTGGAAAGGACGAAACACCG” on the de-multiplexed FASTQ files from Illumina’s NGS sequencer using the Cutadapt software, version 2.8 (https://doi.org/10.14806/ej.17.1.200). FASTQC, version 0.11.9 was used to assess the sequencing quality (http://www.bioinformatics.babraham.ac.uk/projects/fastqc/). MAGeCK, version 0.5.9, was used to quantify and to identify differentially expressed sgRNAs. MAGeCK count command was run on the GECKO library A to quantify. Differentially expressed sgRNAs with statistical significance were determined by running the MAGeCK test command in unpaired mode. Genes were ranked based on the number of unique sgRNA enriched in the mKeima neutral population versus the mKeima acidic population. Data was derived from two biological repeats for each screen.

### Flow cytometry

mKeima-expressing cells were dissociated with fresh DMEM medium by gentle pipetting and passed through a cell strainer cap filter (Thermo Fisher Scientific, catalog no. 08–771-23). Flow cytometry was performed on an LSRII Fortessa analyzer (Becton Dickinson). The gate for acidic (Ex_586_/Em_620_)/neutral (Ex_440_/Em_620_) intensities of individual cells (>10,000 cells) were determined manually using Bafilomycin A1 (100 nM for 2–4 h)-treated samples as a reference. Bafilomycin A1 treatment converts ∼99 % of cell population to the neutral gate. Flow data were analyzed using FlowJo 10.9 software (FlowJo LLC).

### Generation of DNAJC5 antibody

GST-tagged hDNAJC5 purified from E. coli was used for injection to rabbits by a commercial service (LAMPIRE). The GST-DNAJC5 proteins were crosslinked to CNBr-activated sepharose 4 fast flows for antibody purification. In detail, ∼0.85 g of CNBr Activated Sepharose™ 4 Fast Flow (Cytiva, catalog no. 17-0981-01) in 30 mL of ice cold 1 mM HCl was loaded onto a chromatography column (BioRad). The HCl solution is removed by gravity flow and beads were thoroughly washed with 20 mL of ice cold 1 mM HCl three times and 20 mL of coupling buffer (0.1 M NaHCO_3_, 0.5 M NaCl, pH 8.3). GST-hDNAJC5 protein (6 mg) in 15 ml coupling buffer was incubated with the beads for 2 h at room temperature with gentle shaking. The unbound protein was removed by allowing the solution to drain. After washing the beads with 10 mL coupling solution, 20 mL of 0.1 M Tris 8.0 was incubated with the beads for 3 h with shaking at room temperature to block the uncoupled active sites. The buffer was removed and beads were washed 0.1 M glycine pH 3.0, and then with PBS.

To purify antibodies, rabbit anti-DNAJC serum (15 mL) was applied to the column and incubated for overnight with shaking in 4 °C. The antibody was eluted by applying 14 mL of 0.1 M Glycine pH 3.0. The received fractions were combined and applied to Vivaspin 20 centrifugal concentrator (3,000 MWCO; Vivaproducts, catalog no. VS2091) and centrifuged at 4,200 g for 30 min in 4 °C. The resulting solution was dialyzed by PBS with 200 mM NaCl using Slide-A-Lyzer™ Dialysis Cassettes (10K MWCO; Thermo Fisher Scientific, catalog no. 66382) for overnight at 4 °C. The antibody was aliquoted, flash-frozen in liquid nitrogen and stored at −80 °C.

### *Drosophila* experiments

Fly strains bearing shRNA-expressing cassettes downstream of UAS sequences targeting CHIP/STUB1 (33938#), Tsg101 (35710#), Hsp70 (34836#) are from the Bloomington Drosophila Stock Center (BDSC). The flies expressing *human DNAJC5 L116Δ* were described previously ^27^. The strain expressing GAL4 under the heat shock promoter was also purchased from BDSC (2077#). All cultures were maintained on BDSC cornmeal food (Lab-express) in 25 °C incubators equipped with a programmable LED light.

Transgenic flies carrying *UAS-dCHIP* WT, *UAS-dCHIP Δ-Ubox*, or UAS-*dCHIP*-ΔTPR transgenes were made by injecting the corresponding plasmids (a generous gift from Dr. Tang, B. and Duan, R. of Central South University, China) into the R9752 line by Rainbowgene. Stable transgenic lines were crossed to the *Sp/Cyo; Tm2/Tm6* double balancer strain to determine which chromosome harbors the transgene. For most experiments, balanced lines were back-crossed to *W^1118^* to obtain homozygous lines without balancer chromosome. *UAS-Keima-dDNAJC5 (Csp1)* flies were made using a similar strategy.

For imaging fly eyes, 10-20 adult flies were fixed in PBS containing 4% formaldehyde for 1 h, rinsed with PBS. The flies were then dehydrated by soaking sequentially in 30%, 50%, 70%, 90%, and 100% ethanol. The flies were air-dried and dried flies were mounted in an Ibidi imaging chamber by Vaseline. Fly eyes were scanned by a Nikon CSU-W1 confocal microscope using Ex_488_/Em_520_ nm. Shown is the maximum projection view of the scanned Z-section images. Alternatively, flies were fixed on a petri dish with Vaseline and then imaged directly with a dissecting microscope equipped with a AmScope MU1803 digital camera.

Dissecting and immunostaining were performed as previously described ^58^. Briefly, imaginal eye discs were dissected from third instar lava in PBS, fixed with 4% paraformaldehyde in PBS for 20 min at room temperature. Fixed discs were washed with PBS four times, permeabilized with a PBS-based staining buffer containing 0.2% Saponin and 5% FBS in PBS. Discs were then incubated with primary antibodies in the staining buffer at 4 °C overnight. The primary antibodies used are FK2 (1:250) and DNAJC5 (1:500). Discs were washed three times with PBS and then stained with corresponding secondary antibody labeled with either Alexa Fluor 488 or 568.

To detect AFSM and protein aggregates in photoreceptor cells, imaginal eye discs were dissected from third instar larva and incubated in PBS with AmyTracker 680 (Ebba biotech) at 2 µg/mL at room temperature for 30 min and then mounted in Sang M3 medium (Sigma-Aldrich, catalog no. S3652) with 5% FBS and 20 % glycerol. AFSM was detected using Ex405 nm/Em480 nm by a Zeiss LSM780 confocal microscope. To detect apoptotic cells, dissected eye discs were stained with acridine orange (Invitrogen, catalog no. A1301) in PBS at a concentration of 1 µg/mL for 5 min. Discs were washed once with PBS and then mounted for imaging immediately.

### Statistics and reproducibility

All statistical analyses were conducted with GraphPad Prism v10. Statistical methods and the number of cells or *Drosophila* eye discs (N) are indicated in figure legends or shown in figures as individual data point. Biological repeats (n) are specified in figure legends. For statistically significant comparisons with P-value larger than 0.0001, we provided the exact P-values in figures. We did not predetermine sample size. The sample sizes are consistent with similar studies reported in the literature. No biological repeat was excluded from the analyses. For individual cell analyses, a few data points outside of 1.5 times the interquartile range were considered as outliers and were excluded. All experiments were repeated at least twice with individual data point labeled in figures. The immunoblotting and flow cytometry data, whenever shown, are representative of similar results from at least two independent biological replicates unless specified in figure legends. For imaging analyses, cells in randomly selected field were analyzed. The researchers were not blinded. For EM study, lysosomes were identified based on their typical morphology and the presence of intraluminal contents. All identified lysosomes are included in the analysis. Data shown are representative of two biological repeats. For CRISPR screens, two biological repeats were performed with each cell line and the data were pooled to identify statistically significant hits. For organelle-based proteomic study, membranes collected from 3 biological repeats were processed simultaneously. For *Drosophila* experiments, sex was not considered as a variable, and larva were dissected without pre-determination of the sex. All iPSC cell lines were derived from a single male individual. Figures were prepared using ImageJ 1.54f, Adobe Photoshop v25.12.1, and Adobe Illustrator 28.7.4.

## Data Availability

CRISPR screen raw data will be deposited to GEO. All proteomics MS raw files have been deposited to the ProteomeXchange Consortium and are available through the MassIVE repository (Identifier: PXD058788). Other data in support of the conclusions is available in either main figures, extended data figures, or supplementary tables.

## Acknowledgements

We thank the Advanced Light Microscope Core at NIDDK for assistance with imaging, NHLBI flow cytometry core for cell sorting, and NHLBI Genomic Core for high-throughput sequence, S. Yun at NIDDK genomic core for analyzing the CRISPR screen data, K. Zinsmaier (U. Arizona) for GMR-L116Δ flies, R. Puertollano (NHLBI) for critical reading of the manuscript. The research is supported by the intramural research program of NIDDK (Y. Ye), of NICHD (J. Bonifacino), of NINDS (M. Ward), of NCATS (W. Zheng), and by an NIH grant R01NS121608 (L. Hao).

## Author contributions

J. Lee, W. Binti Maxli, N. Chin, M. Jarnik, L. Saidi, Y. Xu, and Y. Ye performed the experiments and analyzed the data. J. Zou and W. Zheng provided the knock-in iPSCs, J. Replogle and M. Ward assisted in i^3^Neuron platform set-up, W. Binti Maxli and L. Hao conducted the mass spectrometry analysis, M. Jarnik and B. Juan conducted the EM analysis. J. Lee, M. Jarnik, L. Hao, and Y. Ye wrote the paper. All authors helped edit the manuscript.

