## Supplementary Figures for "CHIP protects lysosomes from CLN4 mutant-induced membrane damages"

Extended Data Figure 1

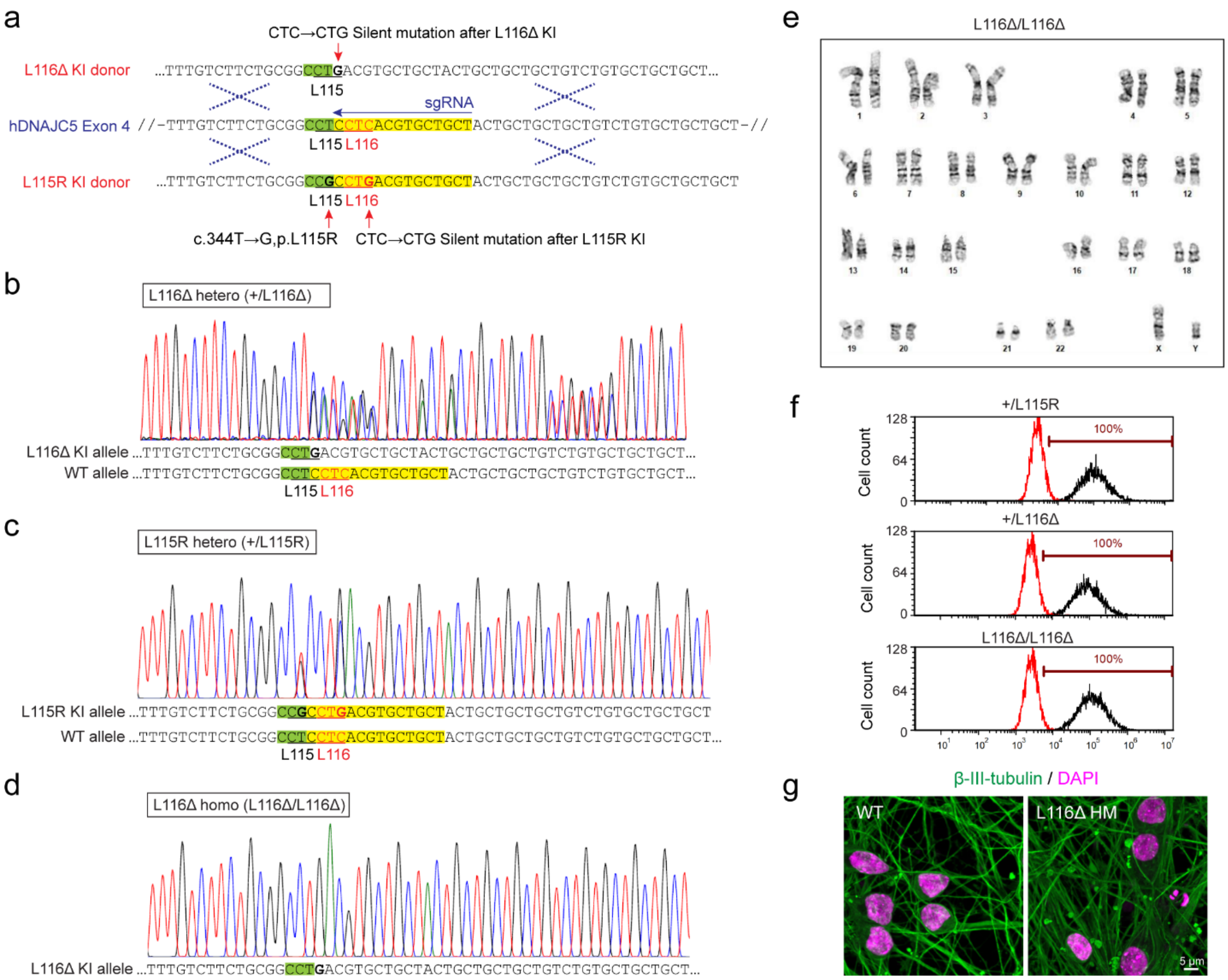


**Extended Data Figure 1. Generation and characterization of human DNAJC5 knock-in (L115R or L116Δ) iPSC lines. a,** Schematic of the gene editing strategy. The codons for L115- and L116 residues are labeled by black and red underlines, respectively. The sgRNA targeting sequence and the following PAM sequence are highlighted in yellow and green. respectively. The knock-in (KI) donor templates with the desired edits (c.344T > G, p.L115R or c.344_346del, p.L116del) and a silent C > G mutation (to avoid recutting by HiFi Cas9 after the knock-in) were shown in parallel to the hDNAJC5 genomic DNA sequence. **b-d,** Sequencing of PCR amplicons confirms the presence either one KI alleles or homozygous L116Δ mutation. **e,** Karyotyping for the HT727D L116Δ homozygous KI iPSC line. The KI line has a normal karyotype with no clonal abnormalities detected at the stated band level of resolution. **f,** Flow cytometry showing the percentage of cells expressing pluripotency markers TRA-1-60 (black peak). The red peaks show cells stained with control IgG. **g,** Wild-type (WT) and L116Δ homozygous (HM) iPSC lines were differentiated as described in the method section. The i$^3$Neurons with indicated genotypes were stained at day16 for a neuronal marker β-III-tubulin (green) and DAPI (magenta). Z-stack images were processed using maximum intensity projection. Scale bars, 5 μm.

Extended Data Figure 2

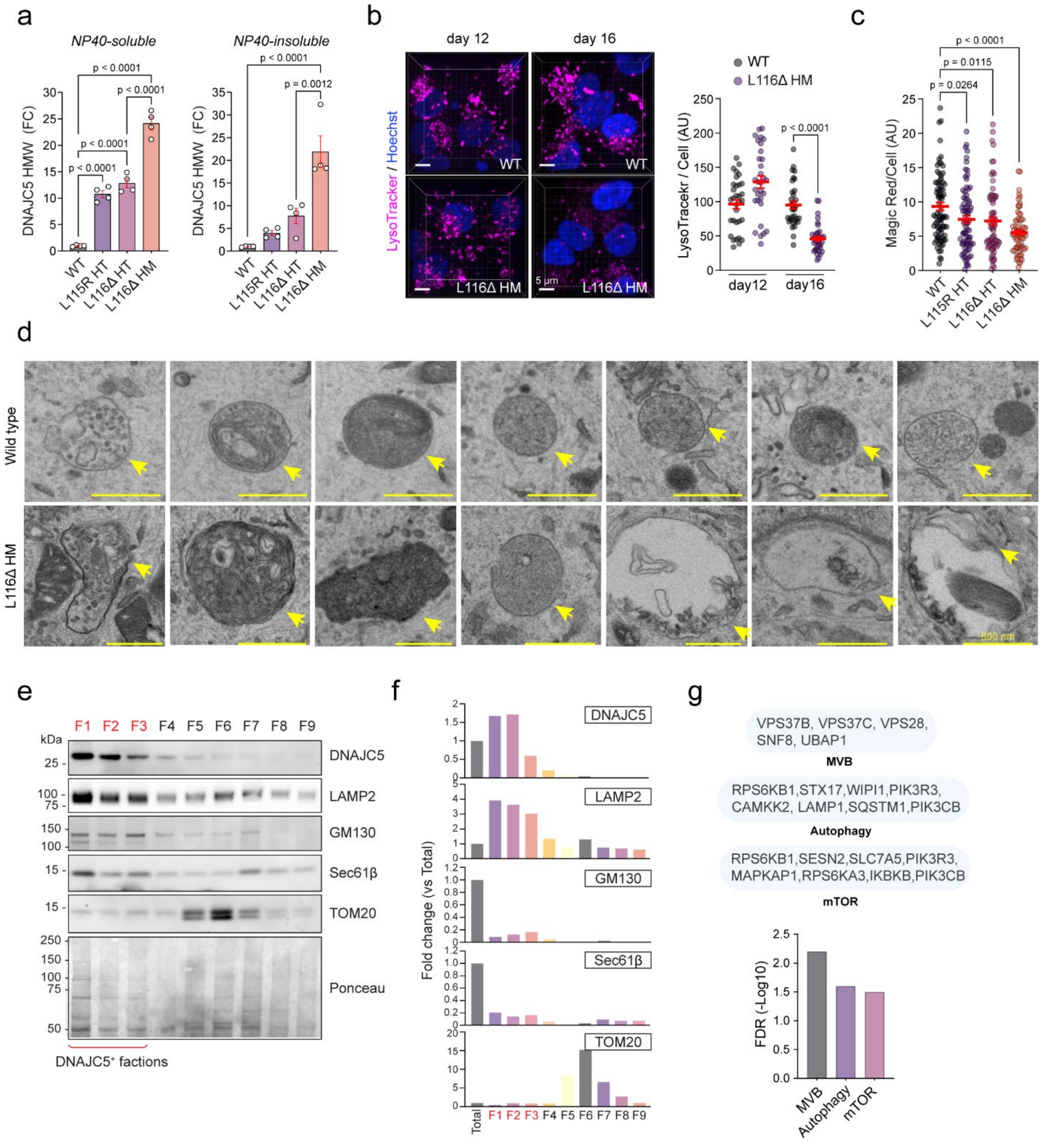


**Extended Data Figure 2. Characterization of lysosomal defects in DNAJC5 L116Δ HM i[3]Neurons. a,** Quantification of the high molecular weight (HMW, >100 kDa) DNAJC5 aggregates in NP40-soluble (left) or insoluble fractions (right) from the indicated i[3]Neurons at d16. Representative blots are shown in Fig 1a. Error bars represent s.e.m. of three biological repeats. P-values were determined by one-way ANOVA. FC, fold change. **b,** 3D reconstructed views of WT and L116Δ homozygous (HM) i[3]Neurons stained with LysoTracker Red (magenta) and Hoechst (blue) at d12 or d16. Scale bars, 5 µm. The right panel shows the quantification of the LysoT signal in randomly selected soma. P-value by unpaired student's t-test. AU, arbitrary unit. **c,** WT and L116Δ HM i[3]Neurons stained with MagicRed and Hoechst at d16 were imaged. The MagicRed signal in individual cells was measured. P-values are from one-way ANOVA. Error bars in b and c represent mean ± standard error of the mean (s.e.m.) (n>20 cells) from two biological repeats. **d,** Additional examples of representative EM images showing lysosomes (yellow arrows) in WT and L116Δ HM i$^{3}$Neurons at d16. Note that irregular, unclosed, and enlarged lysosomes are frequently found in L116Δ HM i$^{3}$Neurons. Scale bars, 600 nm. **e,** Membrane fractions of WT i$^{3}$Neurons were analyzed by immunoblotting with the following antibodies: DNAJC5, LAMP2 (lysosome), GM130 (*cis*-Golgi), Sec61β (ER) and TOM20 (mitochondria). DNAJC5 and LAMP2-enriched fractions (F1-F3 labeled in red) were used for the mass spectrometry analysis. **f,** Quantification of each protein levels in e. e and f represent three biological repeats. **g,** Mass spectrometry analysis of the i$^{3}$Neurons membrane samples showing differentially upregulated proteins (L116Δ vs. WT) in the indicated pathways. The bar graph shows the false discovery rate (FDR, -log10) values for each pathway, indicating their relative significance in the enriched dataset.

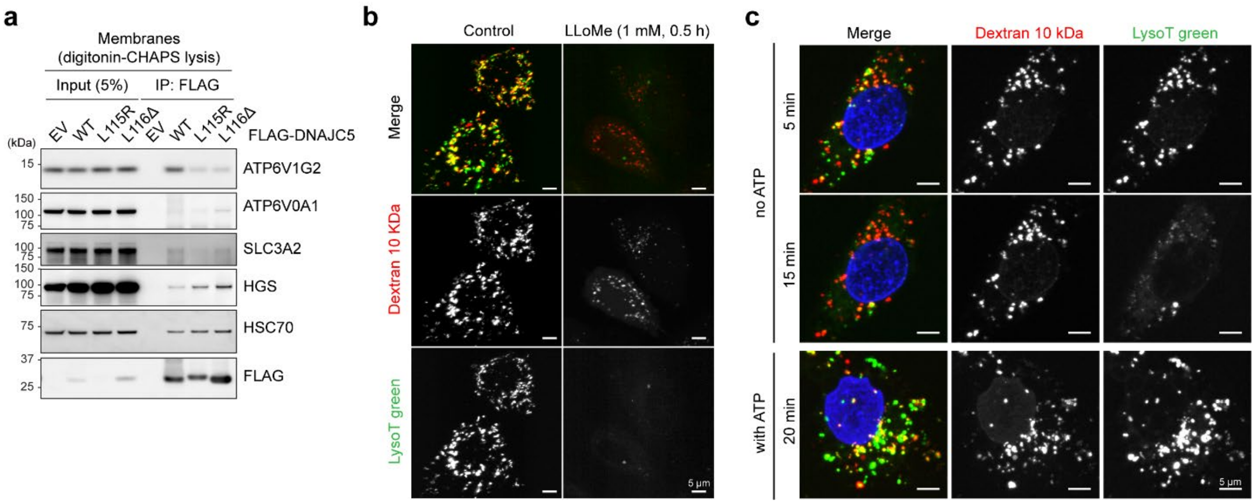


**Extended Data Figure 3. Validation of In vitro lysosome-damaging assay. a,** Interaction between v-ATPase subunits and DNAJC5. HEK293T were transfected with FLAG-tagged DNAJC5 variants as indicated and semi-permeabilized by 0.025% digitonin. The membrane fractions were collected after centrifugation and further lysed by 1% CHAPS lysis buffer as described in the method section. After immunoprecipitation with FLAG M2 agarose, the DNAJC5-bound proteins were analyzed by immunoblotting using the indicated antibodies. SLC3A2 (CD98hc) is a known interactor of DNAJC5 and used as a positive control [1]. **b,** U2OS cells were loaded with Texas Red ™-labeled Dextran (10 kDa) for 4 h, and then incubated in a Dextran-free medium for 3 h. Cells were then labeled by LysoTracker Green dye (1 µM) for 30 min before LLOMe treatment (1 mM, 30 min). Scale bars, 5 µm. **c,** Dextran (10 kDa)-loaded and LysoTracker Green (1 µM, 30min)-stained U2OS cells were permeabilized by SLO in the presence of NucSpot (blue) to distinguish PM-permeabilized cells. Cells were then incubated with a buffer with or without ATP for the indicated time before imaging. In b and c, z-stack images were processed using maximum intensity projection. Scale bars, 5 µm.

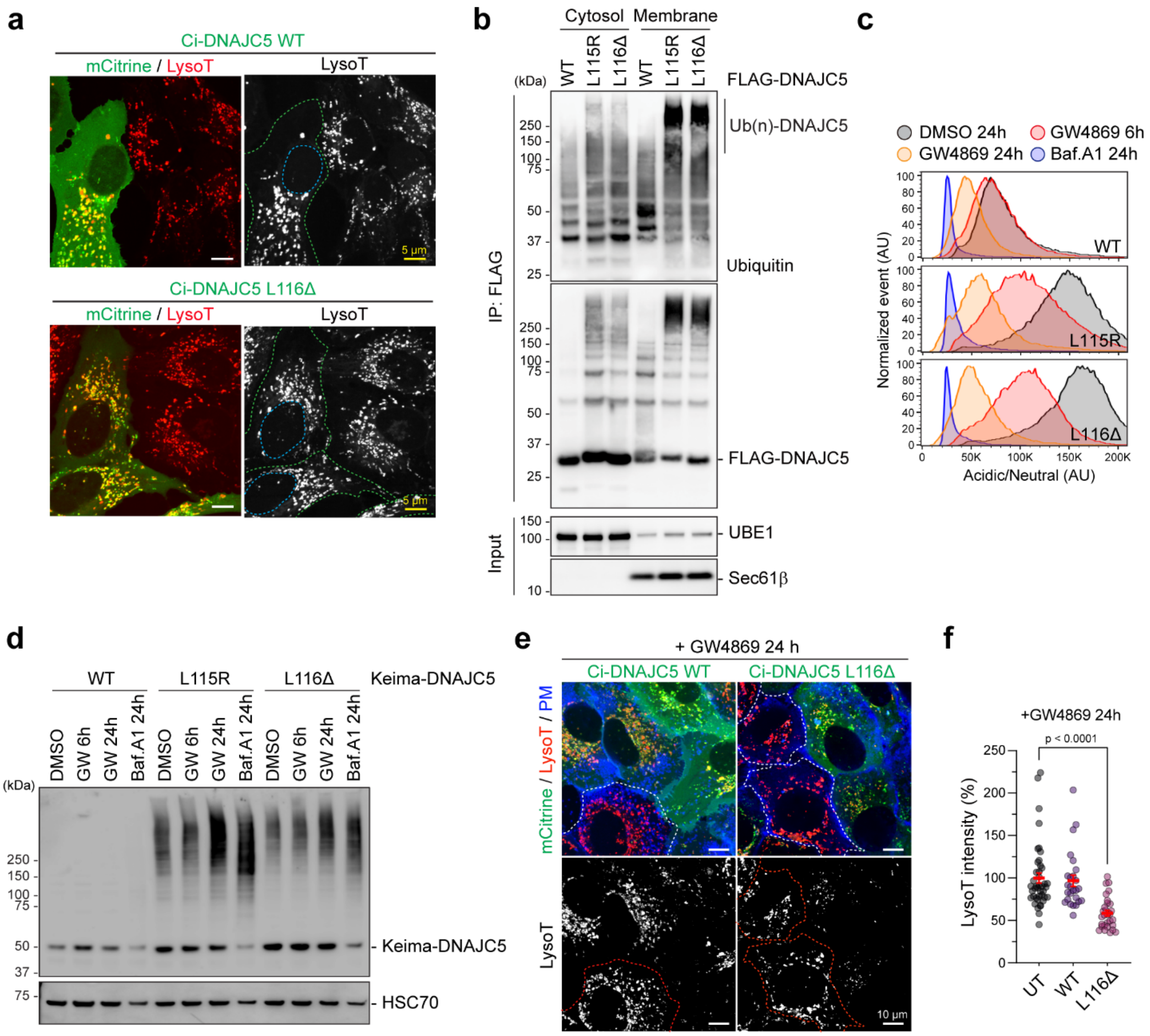


**Extended Data Figure 4. Microautophagy protects lysosomes from CLN4-induced membrane damage. a,** CLN4 mutants does not induce lysosomal defect in U2OS cell. U2OS cells expressing mCitrine (Ci)-tagged DNAJC5 WT or L116Δ were stained with LysoTracker Red and imaged. The green- and blue-dotted lined indicate transfected cells and nuclei, respectively. Scale bars, 5 µm. **b,** Ubiquitinated CLN4 mutants are enriched in cellular membranes. HEK293T cells transfected with the indicated FLAG-DNAJC5 variants were fractionated into cytosol and membrane fractions as described in the method. The fractions were denatured with SDS and the extracts were subject to immunoprecipitation by FLAG beads and immunoprecipitated proteins were analyzed by immunoblotting using the indicated antibodies. UBE1 and Sec61β were as the cytosol and ER marker, respectively. **c,** HEK293T cells stably expressing Keima-DNAJC5 or the indicated CLN4 mutants were treated with GW4969 (10 µM) for 6 and 24 h and analyzed by flow cytometry to measure the acidic/neutral fluorescence ratio of Keima. Bafilomycin A1 (Baf. A1, 20 nM, for 24 h) was used as a positive control. **d,** The Keima-DNAJC5 stable cells used in d were lysed in the NP40 lysis buffer and analyzed by immunoblotting by Keima and HSC70 antibodies. Note that Baf.A1 treatment increased aggregated CLN4 mutants, indicating lysosomal degradation of these species.

**e,** U2OS cells transfected with Ci-DNAJC5 WT or Ci-L116Δ were treated with 10 μM GW4869 for 24 h before staining with LysoTracker (red, top panel) and a PM dye (blue). Untransfected cells were highlighted in the dotted lines. Shown are images of maximum projected view of confocal sections. Scale bars, 10 μm. **f,** Quantification of the relative LysoTracker signal in individual cells as represented in e. P-values were determined by one-way ANOVA from two biological repeats. Error bars represent means ± s.e.m. In a and e, z-stack images were processed using maximum intensity projection.

Extended Data Figure 5

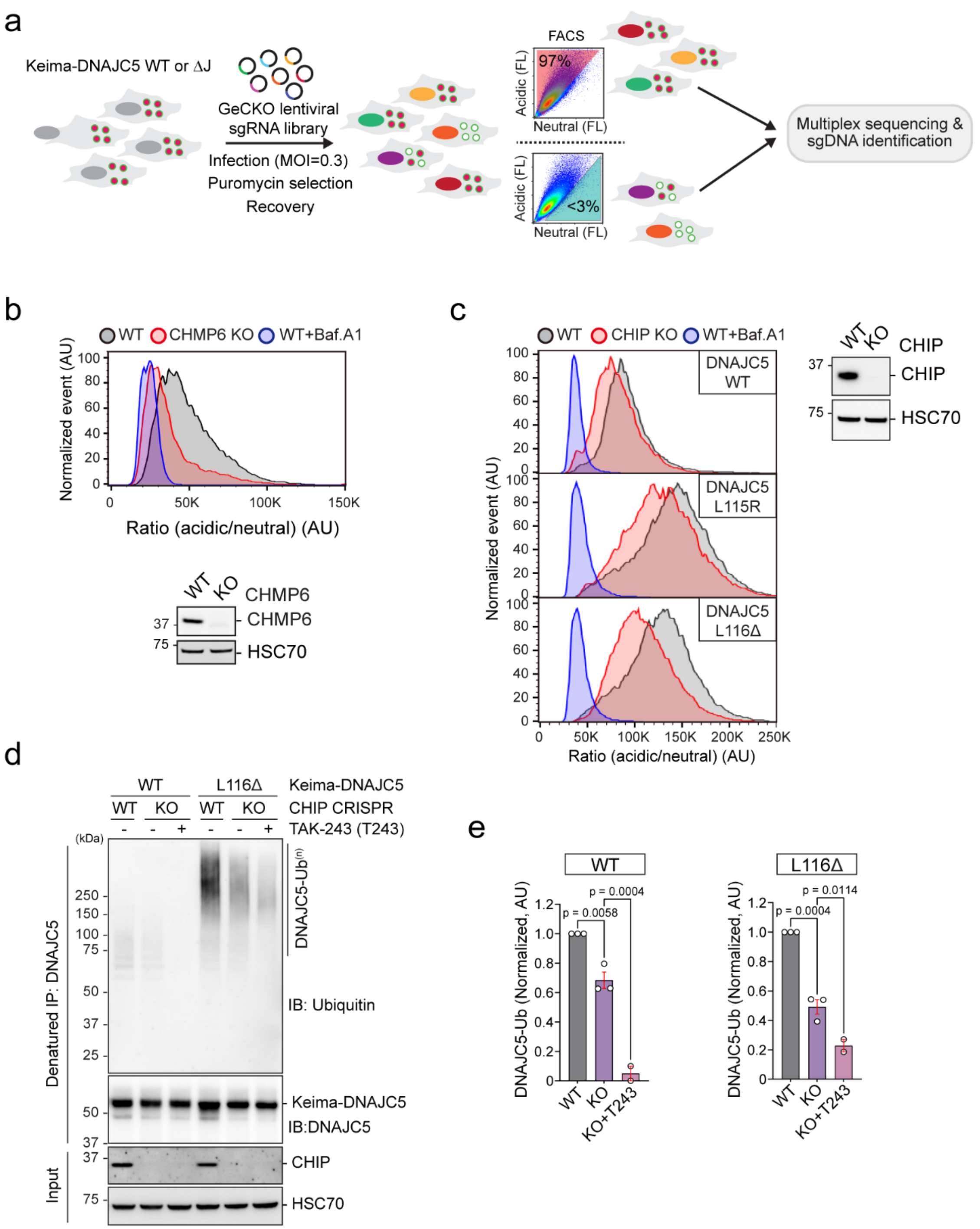


**Extended Data Figure 5. CRISPR screens identified CHIP ubiquitin ligase as a microautophagy regulator. a,** A schematic illustration of the CRISPR screening strategy. **b,** WT or CHMP6 knockout (KO) HEK293T cells were transfected with Keima-DNAJC5 and analyzed by flow cytometry (left) and immunoblotting (right). **c,** CHIP deficiency reduces microautophagy of DNAJC5 WT and CLN4 mutants.

WT or CHIP KO HEK293T cells were infected with lentiviruses expressing the indicated Keima-tagged DNAJC5 variants to obtain stable cell lines, which were then analyzed by flow cytometry (left) and immunoblotting (right). Bafilomycin A1 (50 nM, 4 h)-treated WT cells were used in b and c as controls. **d,** WT or CHIP KO cells infected with Keima-DNAJC5 WT- or Keima-DNAJC5 L116Δ-expressing lentiviruses were drug-selected. Where indicated, stable cells were treated with TAK-243 (1 μM, 4 h). Denatured cell extracts were subject to immunoprecipitation with DNAJC5 antibodies. Precipitated proteins and a fraction of cell lysate were analyzed by immunoblotting. **e,** Quantification of experiments represented by d, Error bars represent means ± s.e.m. of three biological repeats. P values were from one-way ANOVA. T243, TAK-243.

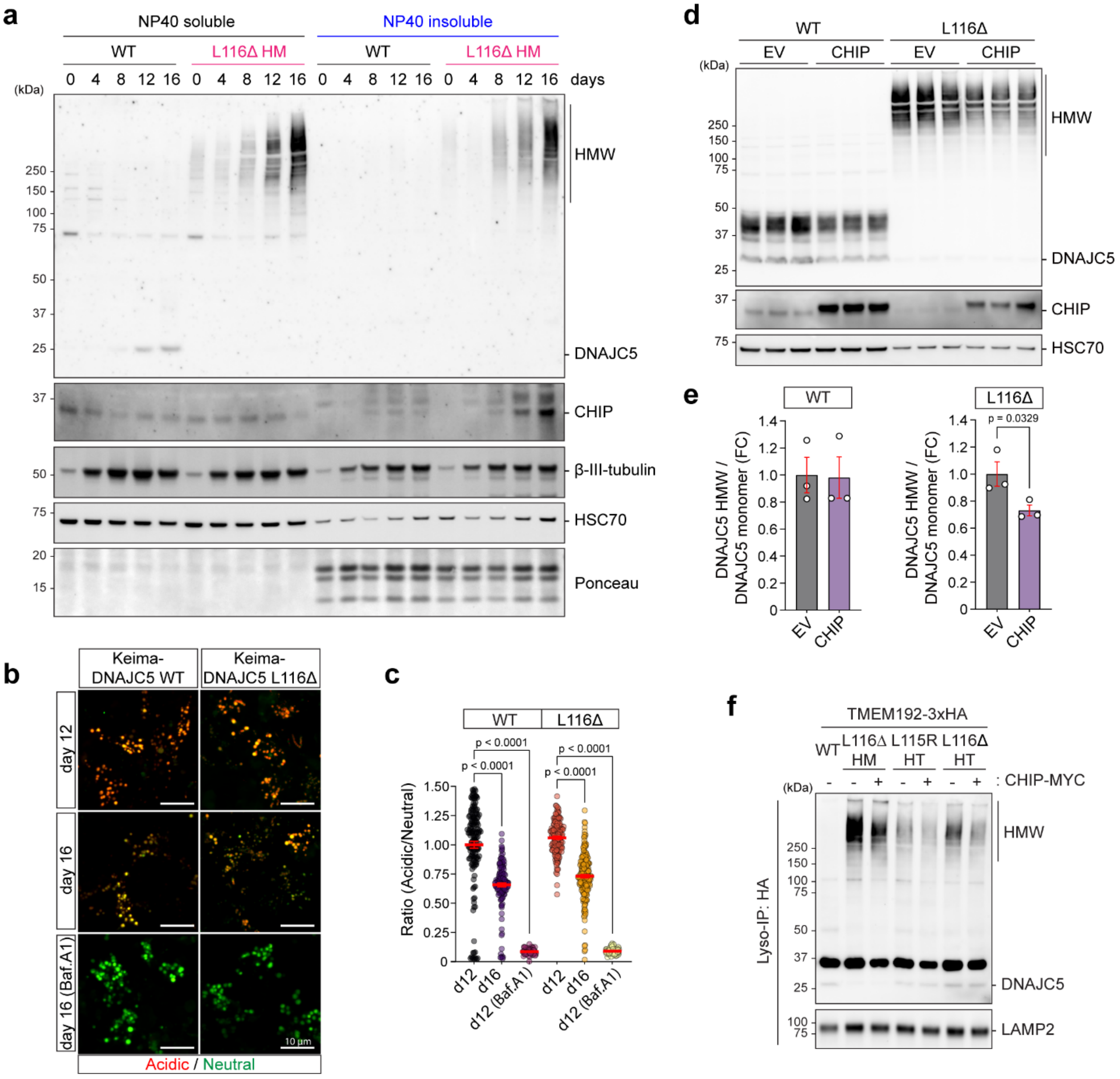


**Extended Data Figure 6. Ectopic expression of CHIP in L116Δ HM i[3]Neurons downregulate CLN4 on lysosomes. a,** A representative blot for Figure 5a. The WT and L116Δ HM i[3]Neuron at indicated days in differentiation were fractionated into NP40 soluble- and insoluble fractions. The samples were analyzed by immunoblotting by the indicated antibodies. Note that CHIP became NP40-insoluble after day12 in L116Δ HM i[3]Neuron. **b,** WT iPSC infected with lentiviruses expressing Keima-DNAJC5 WT or Keima-L116Δ under the Synapsin promoter were differentiated and imaged by confocal microscopy at day 12 and day 16. Baf. A1 treated cells were as a control. z-stack images were processed using maximum intensity projection. Scale bars, 10 µm. **c,** Quantification of acidic/neutral Keima signal ratio in b. Mean intensities from region of interest (ROIs) in each channel were subtract with background before calculating the ratio. P-value are determined by one-way ANOVA. Error bars represent means ± s.e.m. of individual cells (n>50) from two biological repeats. **d,** WT or L116Δ KI iPSC was infected with lentivirus carrying either an empty vector or a Synapsin-controlled CHIP-Myc-expressing cassette before differentiation. The resulting i3Neurons at d16 were lysed by the NP40 lysis buffer and the lysates were analyzed by immunoblotting. Shown are three biological repeats for each condition. **e,** Quantification of the DNAJC5 WT monomer (left) and the high molecular weight L116Δ species in d. P-value is by unpaired student's t-test. Error bars represent means ± s.e.m. of three biological repeats. FC, fold change. **f,** i$^3$Neurons with the indicated genotypes were infected with lentiviruses expressing TMEM192-3xHA and CHIP-Myc (black circles) or TMEM192-3xHA alone (empty circles) at day 6. Lysosomes were isolated at day 16 using the Lyso-IP method as described in the method and then analyzed by immunoblotting using DNAJC5 and LAMP2 antibodies.

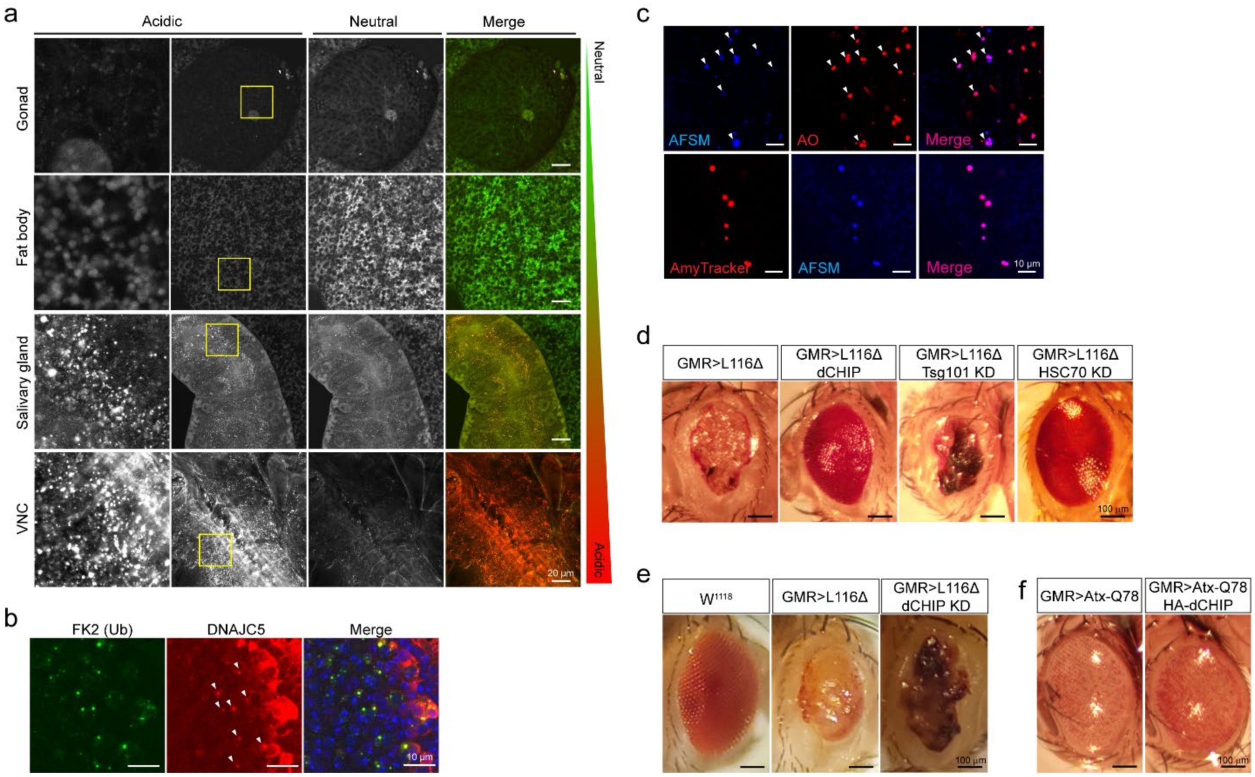


**Extended Data Figure 7. CHIP rescues CLN4 L116Δ-induced neurodegeneration via ESCRT-dependent microautophagy. a,** Third insta larvae of Hs>Keima-dDNAJC5 (*Csp1*) were two rounds of heat shock at 37 degree for 30 min each follow by 30 min recovery. After 24 h, various tissues were dissected and imaged by confocal microscopy. Shown are maximum projected views of confocal sections of the entire tissues. The left panels show enlarged view of the boxed areas. Scale bar, 20 µm. **b,** Imaginal eye discs dissected from third instar larvae of GMR>L116Δ flies were stained with ubiquitin (FK2, green) and DNAJC5 (red) antibodies. Shown are images from a single confocal section. Scale bars, 10 µm. **c,** Imaginal eye discs dissected from third instar larvae of GMR>L116Δ flies were stained with acridine orange (top panels, red) or an Amytracker dye (bottom panels, red), and imaged by confocal microscopy. Arrowheads indicate autofluorescence storage material detected at Ex405/Em450nm. Scale bars, 10 µm. Shown are images from a single confocal section. In a, b, and c, z-stack images were processed using maximum intensity projection. **d,** Adult eyes from flies of the indicated genotypes were imaged. Scale bars, 100 µm. **e, f,** As in d. Scale bars, 10 µm.
